# A surviving beta cell subpopulation enriched in patients with T1D

**DOI:** 10.64898/2026.05.15.725449

**Authors:** Maxwell Spurrell, Zuzana Tobiasova, Jordan S. Pober, John S. Tsang, Kevan C Herold

**Author notes:** **Direct correspondence to:** Kevan C. Herold, MD Department of Immunobiology, Yale School of Medicine, 300 George St, #353E, New Haven, CT 06520.

## Abstract

Type 1 diabetes (T1D) is characterized by the autoimmune destruction of pancreatic beta cells. While most beta cells are lost, a subset of beta cells persists years and even decades after disease onset. Studying these surviving cells is challenging, and thus how they escape immune killing remains poorly understood. Here, we applied a gene regulatory network inference-based clustering approach on existing islet scRNAseq data from cadaveric donors with T1D, autoantibody positive donors at risk for T1D, and non-diabetic donors to analyze beta cells from patients with established T1D. This approach identified a novel beta cell subtype enriched in T1D donors defined by the activity of several transcription factors which have well-characterized roles in beta cell survival, most notably IRF1. We found increased expression of immunomodulatory genes (e.g. *SOCS1/3, HLA-E*) as well as decreased expression of autoantigens and secretory genes, suggesting loss of cellular identity. We identified inflammatory cytokines as a driver of this phenotype by reanalyzing public data from primary human beta cells stimulated with inflammatory cytokines in vitro. We find a similar transcriptional program active in a subset of alpha cells, consistent with cell-extrinsic inflammatory cytokine signaling in vivo. We additionally use immunofluorescence to validate these findings. Overall, this study defines the features of surviving beta cells in patients with T1D, revealing several genes and pathways active in these cells as potential targets for T1D prevention and reversal.

## Introduction

Type 1 diabetes (T1D) is characterized by the autoimmune destruction of insulin-secreting pancreatic beta cells, causing lifelong insulin deficiency (Herold et al., 2024). While most beta cells have been eliminated by the time of T1D diagnosis, a subset of beta cells persists. Even 50 years after initial diagnosis, post-mortem pancreatic tissue examination identified residual beta cells in all assessed T1D patients (Yu et al., 2019). Similarly, residual C-peptide is often found in living T1D patients. Up to 67.4% of participants in the Joslin Medalist study (> 50 years of T1D) had random C-peptide levels in the minimal (0.03-0.2 nmol/L) or sustained range (≥ 0.2 nmol/L), indicating a population of functional, surviving beta cells (Keenan et al., 2010). The reasons why some beta cells persist despite the presence of a primed immune response and autoreactive effector T cells remain poorly understood. Understanding beta cell resilience is highly relevant both for efforts to prevent the disease by protecting existing beta cells and to reverse it with cell replacement therapy. Toward this end, we previously identified a population of resilient beta cells in the non-obese diabetic (NOD) mouse model (Rui et al., 2017). These cells showed reduced granularity by flow cytometry and were found to arise during progressive insulitis, becoming partially dedifferentiated and importantly resistant to both cytokine-induced and T cell-mediated killing.

Less is known about surviving beta cells in humans with T1D since pancreatic biopsies are not routinely performed in living patients. Therefore, persisting beta cells can only be studied from recently deceased cadaveric donors with T1D. The Human Pancreas Analysis Program (HPAP) has performed single cell RNA sequencing (scRNAseq) of islets isolated from donors with T1D, as well as non-diabetic autoantibody positive donors who are at risk for progression to T1D (AAb+) and non-diabetic controls (ND) (Elgamal et al., 2023; Fasolino et al., 2022;

Kaestner et al., 2019; Patil et al., 2023; Ziegler et al., 2013). In this study, we analyzed the public HPAP scRNAseq dataset and focused on the composition and state of beta cells that had survived in donors with T1D. We inferred a gene regulatory network in beta cells and used it to identify a novel population of surviving beta cells through unbiased methods. These cells are characterized by increased activity of known pro-survival transcription factors and decreased transcription of autoantigens and insulin secretory genes. We further show that a similar beta cell phenotype is induced in primary human islets after in vitro cytokine stimulation. Overall, these findings reveal a transcriptional state enriched among surviving beta cells in those with T1D, identifying possible therapeutic targets to promote beta cell survival.

## Results

### Isolation of human beta cells from ND, AAb+ and T1D donors

Islet scRNAseq data from T1D (n = 12), AAb+ (n = 11), or ND (n = 33) donors was downloaded from PANC-DB (full clinical characteristics and assay metadata are in Supplementary Table 1). Among AAb+ donors, 9 had one AAb and 2 had two or more AAbs. After read alignment, initial quality control included removal of cells with few genes detected or high mitochondrial content, correction of ambient transcripts, and automated doublet removal (Methods). We noted that differences in 10X Chromium assays between samples affected the comparability of the data (Supplementary Fig. 1). To correct this, islet cells were integrated using canonical correlation analysis (CCA) (Butler et al., 2018). Clustering of this integrated dataset identified expected cell populations, including endocrine, exocrine, stromal, and immune cell populations (Fig. 1A, Supplementary Fig. 2A-G). We focused on beta cells as they are the primary target in T1D. Beta cells were defined by known secretory products such as insulin (*INS*) and amylin (*IAPP*), as well as the transcription factor *MAFA* (Fig. 1B-C, Supplementary Fig. 2H-I, Supplementary Table 2). To further refine the beta cell population, we performed manual doublet filtering to remove cells expressing known marker genes for other cell populations (Melton et al., 2025) as well as clustering on gene module scores to remove other cells with minimal *INS* expression and expression of known exocrine markers (e.g. *REG1A* and *CFTR*; Supplementary Fig. 3A-G, Supplementary Table 3, Methods). After these refinement steps, we identified 35165 beta cells across all donors (2390 from T1D donors). Importantly, while beta cells were less frequent in T1D samples compared to ND and AAb+ samples, we were able to identify beta cells in nearly all 12 T1D samples (samples HPAP-023 and HPAP-032 had one and zero beta cells respectively and were excluded from downstream analyses; Fig. 1D-E).

**Fig. 1:**
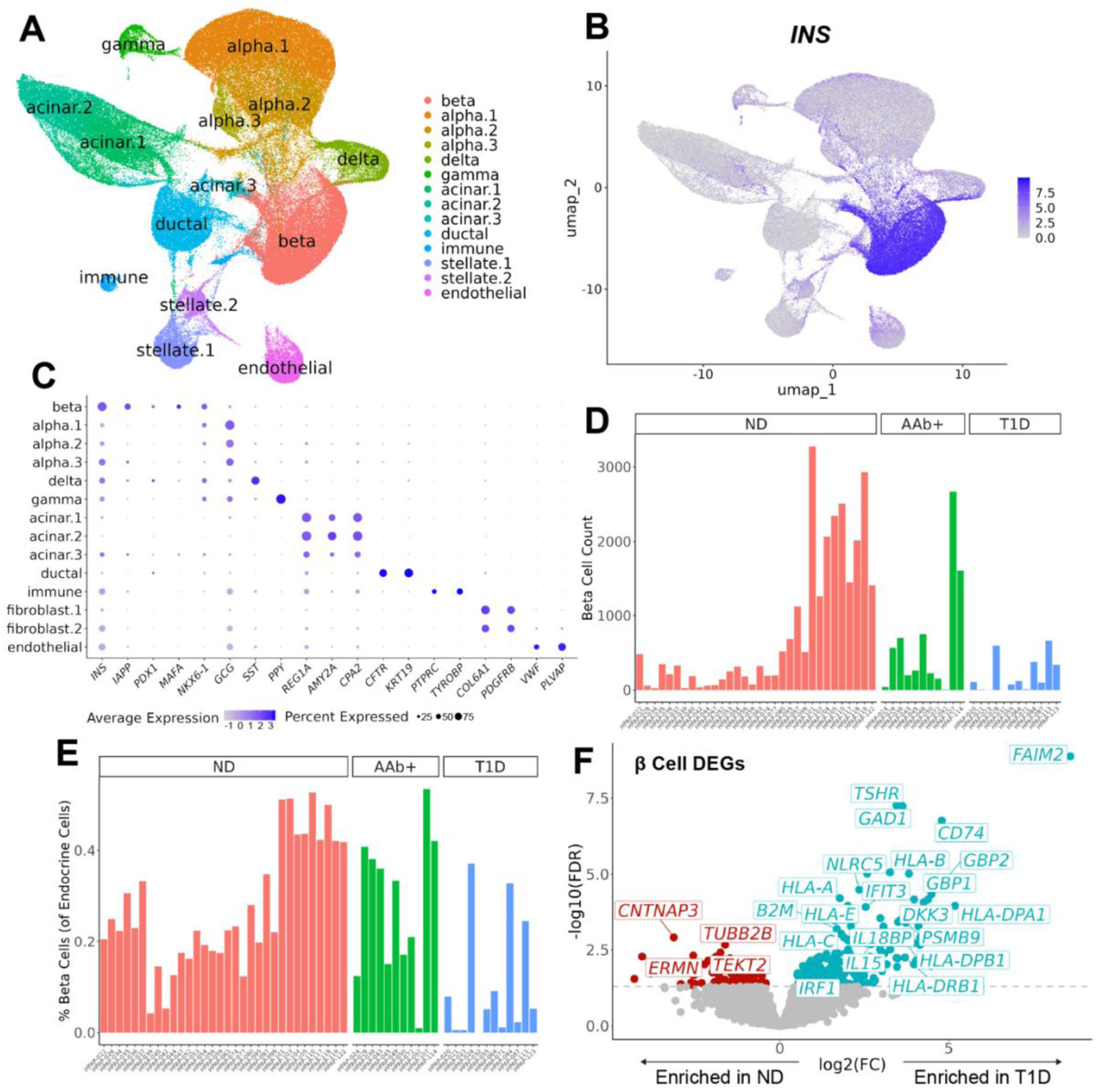
Identification of beta cells in T1D, AAb+, and ND donors from HPAP scRNAseq data. A: Integrated UMAP of islet cell types recovered from HPAP scRNAseq profiling of T1D, AAb+, and ND donors. B: Feature plot showing SoupX-corrected and normalized *INS* expression among islet cells. C: Dot plot of scaled marker gene expression among islet cell populations. A full table of DEGs by islet cell type is available in Supplementary Table 2. D: Beta cell counts per donor. E: Frequency of beta cells per donor as a fraction of total endocrine cells (beta cell count divided by the sum of alpha, beta, delta and gamma cells per donor). F: Volcano plot of DEGs between T1D and ND beta cells calculated using edgeR with donor age, sex, BMI and 10X Chromium kit chemistry as covariates. A pseudobulk method was used by aggregating counts in beta cells by donor (Methods). The Benjamini-Hochberg (BH) procedure was used to calculate FDR. The dashed line indicates FDR = 0.05.

We next tested for differentially expressed genes (DEGs) on the entire refined beta cell population between T1D and ND donors using a pseudobulk approach to account for donor-specific and technical differences (Fig. 1F, Supplementary Table 4, Methods). This revealed that T1D beta cells are enriched for increased class I MHC (e.g. *HLA-A*, *HLA-B*, *HLA-C*, *HLA-E*, *B2M*), class II MHC (e.g. *CD74*, *HLA-DPA1*, *HLA-DPB1*, *HLA-DRB1*), and other interferon-stimulated genes (e.g. *GBP1*, *GBP2*, *NLRC5*, *PSMB9*) versus ND beta cells (434 DEGs with FDR < 0.05). Similar changes, such as increased MHC transcripts, were observed in T1D versus AAb+ beta cells (590 DEGs with FDR < 0.05, Supplementary Fig. 3H, Supplementary Table 4). We did not observe any DEGs between AAb+ and ND beta cells (Supplementary Table 4).

### Identification of a T1D-enriched beta cell subtype through gene regulatory network (GRN) inference

We next sought to identify potential beta cell subpopulations that might be enriched in T1D donors. Distinguishing technical versus true biological heterogeneity is a key challenge in scRNAseq data analysis. While scRNAseq integration methods can correct technical variability (as in Supplementary Fig. 1), integration approaches may overcorrect and lose subtle biological signal present in the uncorrected data (Luecken et al., 2022). Instead, we performed beta cell subclustering on regulon scores calculated using SCENIC, a pipeline for GRN inference using transcriptomic data (Aibar et al., 2017; Van de Sande et al., 2020). This method for GRN inference involves scoring transcription factor (TF) activity using a rank-based approach, which are often robust to technical artefact (Andreatta & Carmona, 2021; Mei et al., 2021; Theodoris et al., 2023). It identifies co-expression between TFs and genes, using known genomic TF binding motif locations to infer direct regulation. A set of genes regulated by each TF (termed a “regulon”) is identified and then regulon activity for a given TF is scored in cells based on the expression level of those target genes. We employed RegDiffusion, a diffusion probabilistic model which artificially introduces noise to identify stable relationships, for the first step of SCENIC GRN inference (Methods) (Zhu & Slonim, 2024). Transcription factors predicted to be active in beta cells along with their target genes are described in Supplementary Table 5. Prior to looking at beta cell subpopulations, we first tested regulons for differential activity between disease states among all beta cells using a pseudobulk-like approach (Fig. 2A, Supplementary Table 6, Methods). This identified that several transcriptions factors were predicted to be more active in beta cells from T1D donors versus ND, including THRB, IRF1, and BCL6 (22 regulons increased and 2 decreased with FDR < 0.05 in T1D). THRB binding motifs have previously been found to be differentially accessible in T1D beta cells, supporting the inference approach adopted here (Melton et al., 2025). T1D beta cells showed similar differences in regulon activity compared to AAb+ beta cells (21 regulons increased and 2 decreased with FDR < 0.05 in T1D, Supplementary Fig. 4A), whereas no differences were observed between AAb+ and ND beta cells (Supplementary Table 6).

**Fig. 2:**
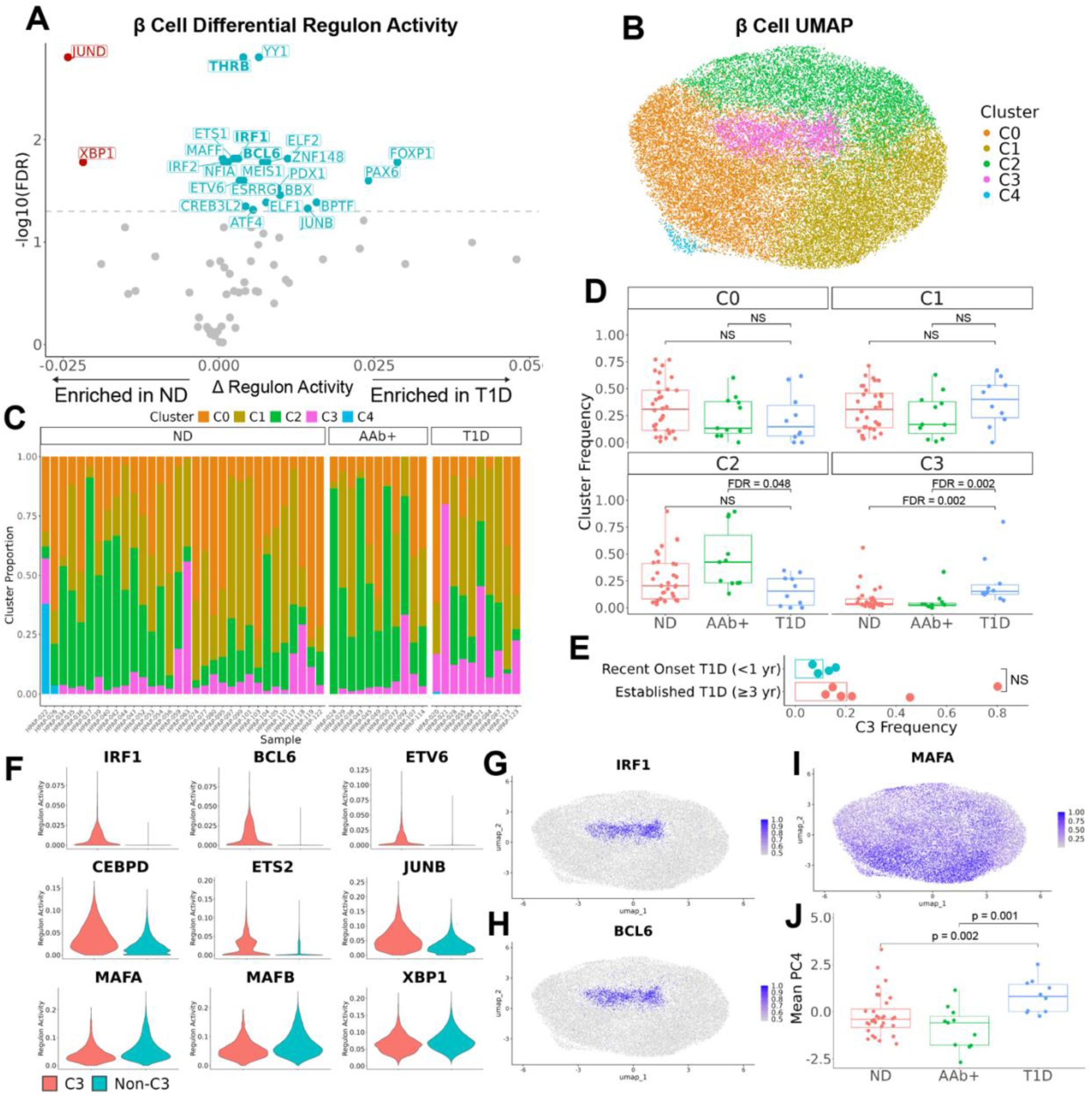
Identification of a T1D enriched beta cell population through GRN inference-based clustering. A: Volcano plot of differentially active regulons calculated using pySCENIC with RegDiffusion (Methods) between T1D and ND beta cells. Regulon scores were averaged by donor. Differential activity testing was performed using a linear model with donor age, sex, BMI and 10X Chromium kit chemistry as covariates. BH-calculated FDR values are shown. The dashed line indicates FDR = 0.05. B: UMAP of beta cells clustered on regulon scores. C: Frequency of each beta cell cluster by donor. D: Frequency of each beta cell cluster by clinical status. Each point is one donor. Disease categories were compared using the Wilcoxon Rank Sum Test. BH-calculated FDR values are shown. NS: not significant. Cluster 4 was not calculated since cells were mainly from a single donor (HPAP-022). E: Frequency of C3 beta cells among donors with recent-onset T1D (less than one year duration) and established T1D (three or more years duration). Each point is one donor. Bars show median values. A Wilcoxon Rank Sum Test was used (p = 0.067). NS: not significant. F: Violin plots of non-rank normalized regulon activity scores between cluster 3 (red) and non-cluster 3 (blue) beta cells. All panels are differentially expressed (Methods) with adjusted p values (Bonferroni) < 0.05. G-I: Feature plots of rank-normalized regulon activity scores. J: Principal component 4 score by clinical status. Each point is the average beta cell PC4 score for one donor. Disease categories were compared using the Wilcoxon Rank Sum Test. Uncorrected p values are shown. The top 6 positive and negative PC4 contributing features were CEBPD, ETS2, JUNB, ETV6, IRF1, BCL6 (positive) and MAFA, MAFB, XBP1, CHD2, ZBTB20, JUND (negative).

To identify whether a subcluster of beta cells drove the observed differential regulon activities, we next performed dimensionality reduction and clustering of beta cells using regulon scores. This revealed several beta cell clusters which were represented across samples and disease states (Fig. 2B-C, Supplementary Fig. 4B-E). In particular, we identified a cluster of beta cells (cluster 3 or C3) that was enriched in samples from donors with T1D (median frequency = 0.153) compared to ND donors (median frequency = 0.037, FDR = 0.002 by Wilcoxon Rank Sum test) and AAb+ donors (median frequency = 0.027, FDR = 0.002 by Wilcoxon Rank Sum test) (Fig. 2D). This population of cells was stably detected across several clustering resolutions (Supplementary Fig. 5). Interestingly, the two AAb+ donors with the highest C3 frequencies (HPAP-092 and HPAP-107) have been independently predicted to be most likely to progress to T1D among HPAP AAb+ donors using an XGBoost classifier trained across all cell types (Patil et al., 2024). A small number of ND samples likewise had elevated C3 frequencies which may be a reflection of the older age of certain ND donors (the average age of the T1D donors was 14.9 years—6 of 32 ND donors had greater than 10% of beta cells assigned to C3 and all 6 donors were at least 35 years old; Supplementary Fig. 4F-G). Among T1D donors, the frequency of C3 beta cells was greater in donors with established T1D (median frequency = 0.203) compared to those with recent-onset T1D (median frequency = 0.109), though the difference was not statistically significant (p = 0.067, Wilcoxon Rank Sum test, Fig. 2E).

We performed differential regulon activity testing between clusters to identify which regulons were enriched in cluster 3 (Supplementary Table 7). The TFs with the greatest increase in activity in cluster were IRF1, BCL6, ETV6, CEBPD, ETS2, and JUNB, many of which have previously been reported to promote beta cell survival and modulate beta cell immune responsiveness (Fig. 2F-H, Supplementary Fig. 4H-K) (Baker et al., 2003; Colli et al., 2018; Colli et al., 2020; Gurzov et al., 2012; Gurzov et al., 2008; Gysemans et al., 2009; Igoillo-Esteve et al., 2011; Moore et al., 2011; Moore et al., 2012). Several of these regulons were enriched in T1D when comparing all beta cells between T1D and ND (Fig. 2A), suggesting increased C3 frequencies may be driving those differences. We also identified regulons with decreased activity in C3 including the lineage-specific factors MAFA and MAFB as well as the ER stress regulator XBP1 (Fig. 2F&I, Supplementary Fig. 4L-M) (Lee et al., 2011; Wortham & Sander, 2021).

Given that each of these transcription factors was represented by the fourth principal component (PC4) while performing dimensionality reduction on regulons for clustering, we compared beta cell PC4 scores across disease states to test whether we could detect changes along this axis independent of clustering. Supporting this notion, we found that PC4 was elevated in T1D samples versus both ND samples (p = 0.002 by Wilcoxon Rank Sum test) and AAb+ samples (p = 0.001 by Wilcoxon Rank Sum test, Fig. 2J). We then compared regulon activity within each cluster across disease states (Methods). We did not detect differences between T1D and ND/AAb+ C3 beta cells (Supplementary Fig. 6, Supplementary Table 6), implying that while the frequency of these cells varies by clinical status, the underlying TF activity is similar.

To assess the stability of our SCENIC workflow, we retrained the RegDiffusion model 9 additional times and re-scored regulons, then compared to the initial training. While we observed minor run-to-run differences, the top differentially active regulons in C3 were detected by SCENIC in most runs and showed consistent correlation trends across runs (Supplementary Fig. 7).

### Transcriptional profile of cluster 3 beta cells

Beyond inferred TF activities, we next examined which transcripts distinguished C3 beta cells from other beta cells. The comprehensive list of DEGs for each cluster is found in Supplementary Table 8. The most highly enriched genes for C3 are shown in Fig. 3A, which included genes involved in suppression of cytokine signaling (*SOCS3* and *SOCS1*). Beta cell *SOCS1* and *SOCS3* expression has previously been shown to promote survival after in vitro inflammatory cytokine stimulation and prevent diabetes in murine models (Benaglio et al., 2022; Chong et al., 2004; Flodström-Tullberg et al., 2003; Karlsen et al., 2001; Rønn et al., 2008).

**Fig. 3:**
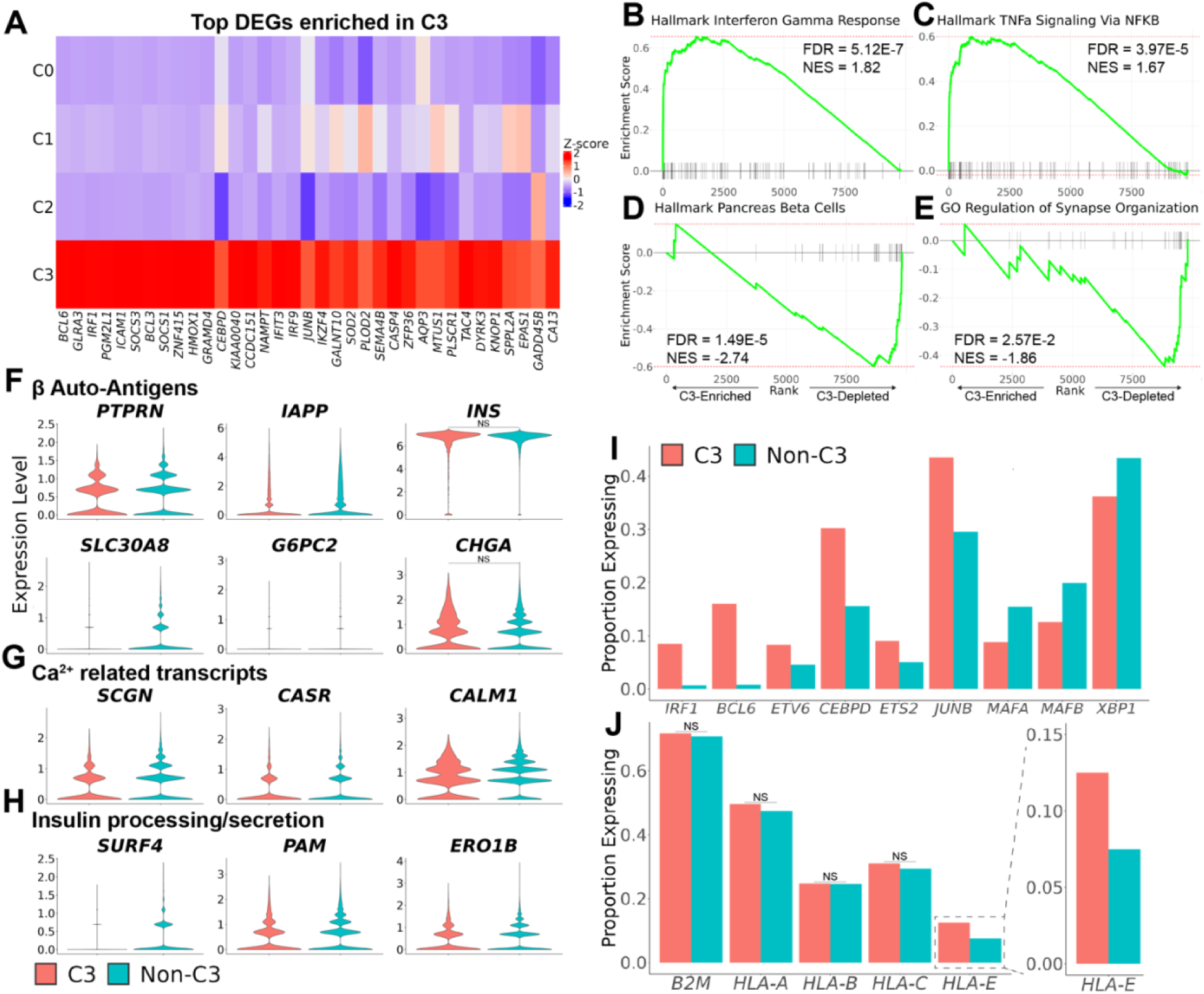
Transcriptional profile of cluster 3 beta cells. A: Heatmap of average expression for each beta cell cluster of the top 35 positive differentially expressed genes in C3 (by adjusted p value). Values were scaled for visualization. Cluster 4 was omitted (prior to scaling) since cells were mainly from a single donor (HPAP-022). B-E: Gene set enrichment plots for selected pathways (full list of gene sets in Supplementary Table 3). Genes were ranked by fold change in C3 beta cells (vs non-C3 beta cells), with filtering of lowly expressed genes (Methods, full table of GSEA output in Supplementary Table 10). BH-calculated FDR values are shown. NES: normalized enrichment score. F-H: Violin plots showing expression of selected transcripts for C3 and non-C3 beta cells. All panels are differentially expressed by Wilcoxon-based testing (Methods) with adjusted p values (Bonferroni) < 0.05 unless indicated by NS (not significant). I-J: Frequency of beta cells expressing TFs (I) and class I HLA genes (J) between C3 and non-C3 beta cells. For J, an inset for *HLA-E* is shown. All adjusted p values (Bonferroni) < 0.05 unless indicated by NS (not significant). P values derived from Wilcoxon-based DEG testing. Full C3 differential expression tabular data is available in Supplementary Table 9.

While interferon stimulated genes (ISGs) were expressed by only a small proportion of beta cells, we found that C3 had increased expression of several ISGs in addition to *SOCS1/3* (*ICAM1*, *NLRC5*, *STAT1*, *IFIT3*, *PSMB8*, *IFITM3*, *CD274*; Supplementary Fig. 8A, Supplementary Table 9). C3 DEGs additionally contained certain stress response genes (*SOD2*, *HMOX1*, *IER3*, *GADD45A/B/G*, *CDKN1A*, *DDIT4*) associated with oxidative stress and/or DNA damage (Supplementary Fig. 8B). We next performed gene set enrichment analysis (GSEA) on genes ranked by C3 differential expression (Supplementary Table 10) (Mootha et al., 2003; Subramanian et al., 2005). Importantly, C3 beta cells were found to be enriched for inflammatory cytokine programs such as the Hallmark “Interferon Gamma Response” (FDR = 5.12E-7) and “TNFa Signaling Via NFKB” pathways (FDR = 3.97E-5, Fig. 3B-C).

Interestingly, negatively enriched pathways in C3 included pathways related to beta cell identity and exocytosis, namely the Hallmark “Pancreas Beta Cell” pathway (FDR = 1.49E-5) and the GO “Regulation of Synapse Organization” pathway (FDR = 2.57E-2, Fig. 3D-E).

Concordantly, we observed decreased expression of a subset of beta cell autoantigens (*IAPP*, *G6PC2*, *SLC30A8*, and *PTPRN*; Fig. 3F, Supplementary Table 9) (Purcell et al., 2019). We did not observe differences in *INS, CHGA*, or *GAD2* expression (Fig. 3F, Supplementary Fig. 8C). We additionally found that C3 exhibited decreased expression of certain transcripts with calcium-related activity *(SCGN*, *CASR*, *CALM1*, *CADPS*) and roles in insulin processing and secretion (*PAM*, *ERO1B, SURF4, SEC24D, VAMP2*; Fig. 3G-H, Supplementary Fig. 8D-E) (Chen et al., 2025; Grenko et al., 2024; Regazzi et al., 1995; Saegusa et al., 2022; Viviano et al., 2020). Proinsulin proteases *PCSK1* and *CPE* (and to a lesser extent *PCSK2*) were similarly expressed at lower levels in C3 beta cells (Supplementary Fig. 8F-H) (Urbaniak et al., 2025).

Overall, these changes suggest that C3 beta cells show reduced expression of genes associated with beta cell identity.

We next assessed the expression of TFs predicted to be differentially active in cluster 3, finding expected increased expression of *IRF1*, *BCL6*, *ETV6*, *CEBPD*, *ETS2*, and *JUNB* and decreased expression of *MAFA*, *MAFB*, and *XBP1* (Fig. 3I). We also examined the expression of other key beta cell identity TFs, such as *NKX6-1*, *PDX1*, *NEUROD1*, and *PAX6*, finding that only *NKX6-1* showed decreased expression in C3, consistent with partial loss of cellular identity (Supplementary Fig. 8I, Supplementary Table 9) (Wortham & Sander, 2021). While we had previously found that class I HLA genes were differentially expressed among all beta cells between T1D and ND/AAb+ (Fig. 1F, Supplementary Fig. 3H), this was not due to increased class I HLA in C3. We observed that only *HLA-E* showed enrichment in C3, with no difference in *B2M* and *HLA-A/B/C* between C3 and other clusters (Fig. 3J). Instead, class I HLA genes were differentially expressed between T1D and ND/AAb+ donors across multiple clusters, suggesting that class I HLA hyperexpression is a more general feature of T1D beta cells (Supplementary Fig. 9, Supplementary Table 4). These results are consistent with reports of a general increase in class I MHC on islet cells from T1D patients (Benkahla et al., 2021; Foulis et al., 1987; Patil et al., 2024).

We then tested whether the transcriptional trends in C3 could be identified within individual T1D donors. We focused on the 3 T1D donors with the greatest number of C3 beta cells (HPAP-028, HPAP-113, and HPAP-123) and compared their transcriptional profile to non-C3 beta cells from the same donors (Supplementary Fig. 8J). While underpowered to test for significance due to the scarcity of beta cells in individual T1D donors, we qualitatively observed similar transcriptional changes between C3 and non-C3 beta cells in these donors. Altogether, these findings demonstrate that C3 beta cells are characterized by inflammatory response genes with decreased production of autoantigens and secretory transcripts.

### Cluster 3-like phenotype is reproducible by stimulation with inflammatory cytokines

To confirm our findings and determine whether inflammatory mediators can induce the changes we observed, we reanalyzed scRNAseq data from Maestas et al. which profiled primary human islets from non-diabetic donors (n = 5) treated in vitro with various stressors, including inflammatory cytokines (IFNG, TNFA, and IL1B; cytokine mix [CM] refers to co-stimulation with all three) or ER stress inducers (Brefeldin A [BFA] and Thapsigargin [TG]; all stressors were applied for 48 hours) (Maestas et al., 2024). After re-processing the data, we identified beta cells representing each donor and treatment combination (Methods, Supplementary Fig. 10A-H, Supplementary Table 11).

We then tested whether any treatments caused similar gene expression changes to those observed in C3. This was performed by scoring pseudobulk transcript counts (aggregated by donor and treatment combination) for gene sets comprising the top 50 up- and down-regulated genes in C3 using GSVA (Methods) (Hänzelmann et al., 2013). Importantly, we observed that all cytokine conditions (either alone or in combination) caused overexpression of C3-enriched transcripts and decreased expression of C3-depleted transcripts (Fig. 4A, Supplementary Table 12, FDR < 0.05 versus control beta cells for all cytokine condition model coefficients). In contrast, ER stressors led to unidirectional changes, with BFA causing downregulation of the C3 down gene set and TG causing upregulation of the C3 up gene set. Stimulation with CM produced the most extensive bidirectional change, suggesting that CM stimulation can produce the most C3-like phenotype. Similar to our findings with the HPAP dataset, we observed that CM-treated beta cells increased expression of *IRF1*, *BCL6*, *ETV6*, *CEBPD*, *ETS2*, and *JUNB*, whose activity defined C3 (Methods, Fig. 4B, Supplementary Table 13). We likewise observed decreased expression of *MAFA* and *MAFB*, which had been found in C3 (Fig. 4B). Interestingly, CM-treated beta cells did not show reduced expression of *XBP1* (predicted to be less active in C3, Fig. 4B). *XBP1* was instead reduced in BFA-treated beta cells (Fig. 4C). We also observed that while CM-treated beta cells expectedly expressed many ISGs, only a subset of these ISGs were also enriched in C3 beta cells (Fig. 4D, Supplementary Fig. 10I, Supplementary Tables 9 & 13). These findings suggest that cytokines reproduce many of the features of the C3 phenotype in beta cells.

**Fig. 4:**
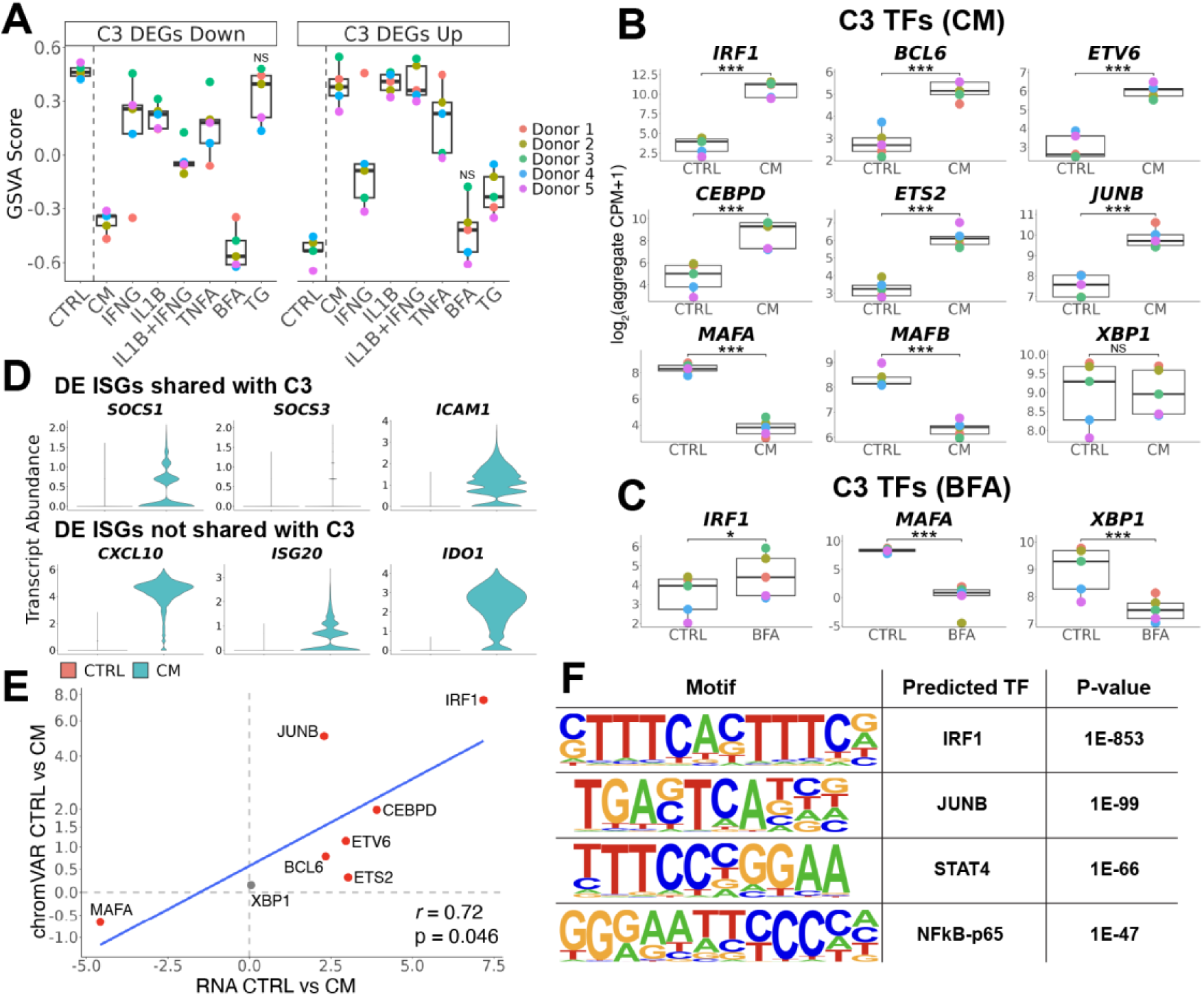
In vitro cytokine stimulated beta cells are similar to cluster 3 beta cells. Beta cells treated in vitro for 48 hours with various stressors were identified from scRNAseq data from Maestas et al. (Methods, Supplementary Fig. 10A-H). A: In vitro beta cell expression of C3 up and down gene sets. Gene sets were synthesized by taking the top 50 non-ribosomal/mitochondrial up- and down-regulated genes (by p value) in HPAP C3 beta cells (Methods). GSVA was used to calculate pathway scores for aggregated (pseudobulk) counts for each sample. Each point is a GSVA score for one sample, colored by donor. CTRL: control, CM: cytokine mix with IFNG+IL1B+TNFA (1000 ng/mL, 100 ng/mL, and 500 ng/mL, respectively), BFA: Brefeldin A (1 μg/mL), TG: Thapsigargin (10 μM). All have BH-calculated FDR < 0.05 versus CTRL unless indicated by NS (not significant). B-C: Paired expression plots for indicated transcripts in beta cells treated with cytokines (B) or BFA (C) versus control beta cells. Log_2_(counts per million + 1) of aggregated (pseudobulk) counts per sample are shown. Each point is the expression for one sample, colored by donor. Differential expression testing was performed using edgeR with donor as a covariate (Methods). BH-calculated FDR values are shown. Full tabular data for Maestas et al. differential gene expression is found in Supplementary Table 13. ***: FDR < 0.001, *: FDR < 0.05, NS: not significant. D: Violin plots of in vitro cytokine-treated and control beta cells (pooled from all donors). All panels are differentially expressed (Methods) with adjusted p values (Bonferroni) < 0.05. Shared ISGs are those differentially expressed in both in vivo C3 vs non-C3 beta cells (HPAP) and in vitro CM-treated vs control beta cells (Maestas et al.). Non-shared ISGs are those only differentially expressed by in vitro CM-treated vs control beta cells (Maestas et al.). Full tabular data for HPAP C3 DEGs are found in Supplementary Table 9 and for Maestas et al. in Supplementary Table 13. E: Changes in beta cell motif accessibility versus transcript abundance between CM and CTRL treatment (48 hour stimulation, same cytokine concentrations as for Maeastas et al. scRNAseq). Beta cells were isolated from Maestas et al. snMultiome data (Supplementary Fig. 11). Differential motif accessibility was calculated using chromVAR (Supplementary Table 15). Difference in mean chromVAR z-score for CM beta cells minus mean CTRL z-score is shown on the y-axis. Log_2_ fold change between CM and CTRL beta cells is shown on the x-axis, calculated from scRNAseq pseudobulk DEG testing (panel B). TFs with significant changes in both transcript abundance and motif accessibility are colored red. Linear regression between differential transcripts and differential motif accessibility (blue line) and Pearson correlation coefficient are shown. F: Top hits of de novo HOMER motif analysis of differentially accessible beta cell snATACseq peaks after IFNG+IL1B+TNFA stimulation (24 hour stimulation with 10 ng/mL, 0.5 ng/mL, and 1 ng/mL, respectively) from Benaglio et al. (Methods).

We also asked whether these TF changes could be detected on the chromatin level after cytokine stimulation, which would support active transcriptional modulation by these TFs. We leveraged single nucleus multiome profiling (RNA + ATAC) of CM and control treated islets (also from Maestas et al., islets from n = 1 ND donor, 48 hour treatment duration). After reprocessing the data and identifying beta cells (Supplementary Fig. 11, Supplementary Table 14), we scored beta cells for TF motif availability using chromVAR (Fornes et al., 2020; Schep et al., 2017). We then tested for differential motif accessibility between CM and control beta cells (Supplementary. Table 15). When comparing to differences in transcript abundance (from Fig. 4B), we observed highly concordant changes in motif accessibility, suggesting functional TF binding of C3 TFs (Fig. 4E, Pearson correlation between RNA and chromVAR = 0.72, p = 0.046, MAFB excluded since it is not annotated in the JASPAR2020 motif database).

We further validated these findings using a third data set from Benaglio et al. of single-nucleus chromatin profiling (snATACseq) of primary human islets from n = 3 ND donors stimulated for 24 hours with a combination of IFNG, IL1B, and TNFA or control (Benaglio et al., 2022). We performed de novo motif analysis using HOMER within differentially accessible chromatin regions between stimulated and unstimulated beta cells (Methods, Supplementary Table 16) (Heinz et al., 2010). The top enriched motif within these peaks was assigned to IRF1 (Fig. 4F). A motif assigned to JUNB was also identified as the second most enriched sequence (Fig. 4F), providing additional support that these TFs are critical for the beta cell response to inflammatory cytokines.

### Homologous alpha cell population supports cytokines as driver of cluster 3 phenotype in vivo

To further test the hypothesis that C3 beta cells are those experiencing cytokine stress, we reasoned that any secreted cytokines should also influence other proximal cell populations in vivo. We therefore focused on developmentally-related alpha cells to test whether: 1) we could observe evidence of cytokine exposure in samples with elevated C3 beta cell frequencies and 2) whether the C3 phenotype is unique to beta cells. We analyzed alpha cells in the HPAP dataset (Fig. 1A) with the same approach used for beta cells (Supplementary Fig. 12), clustering on regulon scores inferred independently in alpha cells (Fig. 5A, Supplementary Fig. 13A-E). This identified an alpha cell population (alpha cell cluster 3) with very similar regulon activity to C3 beta cells. C3 alpha cells had increased IRF1, BCL6, CEBPD, JUNB, and ETV6 regulon activity relative to other alpha cells (Fig. 5B, Supplementary Fig. 13F, Supplementary Table 17). Additionally, MAFB was decreased in C3 alpha cells, consistent with loss of cellular identity (Supplementary Fig. 13G).

**Fig. 5:**
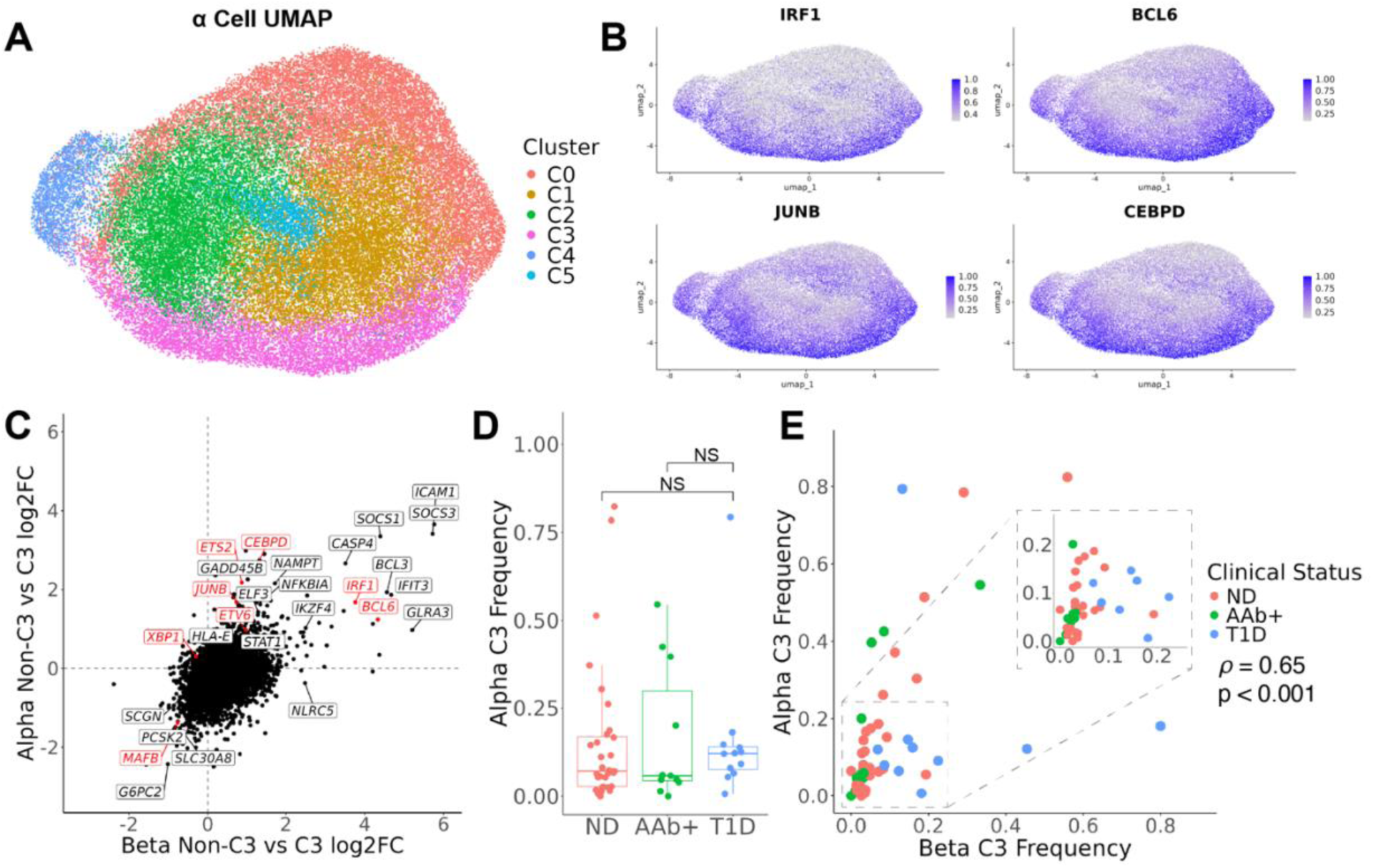
Identification of a homologous alpha cell population to cluster 3 beta cells. A: UMAP of alpha cells from the HPAP dataset clustered on regulon scores. B: Feature plots of rank-normalized regulon activity scores. C: Gene expression log_2_ fold change values for C3 beta cells versus non-C3 beta cells (x axis) plotted against log_2_ fold change values for C3 alpha cells versus non-C3 alpha cells (y axis). Genes were filtered to those expressed by ≥ 1% of cells in at least one test group within both comparisons. Full tabular results are in Supplementary Tables 9&19. Transcripts of regulons used to define C3 beta cells are shown in red. D: Frequency of C3 alpha cells by clinical status. Each point is one donor. Disease categories were compared using the Wilcoxon Rank Sum Test. NS: not significant. E: Frequency of C3 beta cells among all beta cells plotted against frequency of C3 alpha cells among alpha cells for each donor with a boxed inset. The Spearman correlation coefficient (ρ) is shown (p = 1.49E10-7).

We next compared the relative transcriptional changes observed in C3 alpha cells to those in C3 beta cells (Fig. 5C, Supplementary Tables 18 & 19). We observed that C3 alpha cells express many of the same genes as C3 beta cells. *ICAM1*, *SOCS1*, *SOCS3*, *IFIT3*, *HLA-E*, and *STAT1* were elevated in C3 alpha cells. Likewise, we observed concurrent decreases in alpha cell functional genes, including *SCGN*, *CASR*, *G6PC2*, *SLC30A8*, and *PCSK2*. On the pathway level, C3 alpha cells were enriched for the “Interferon Gamma Response” (FDR = 1.20E-15) and “TNFa Signaling Via NFKB” pathways (FDR = 4.63E-31, Supplementary Fig. 13H-I, Supplementary Table 20).

We observed a non-significant trend towards increased C3 alpha cell in donors with T1D (median frequency = 0.12) relative to ND (median frequency = 0.07, p = 0.397, Wilcoxon Rank Sum Test) and AAb+ donors (median frequency = 0.06, p = 0.413, Wilcoxon Rank Sum Test, Fig. 5D). As with C3 beta cells, C3 alpha cell frequency was increased in older ND donors, likely confounding statistical testing (Supplementary Fig. 13J). Importantly, we found a strong positive correlation between C3 beta cell frequency and C3 alpha cell frequency (Spearman’s ρ = 0.65, p = 1.49E10-7), supporting the notion that C3 beta cells and C3 alpha cells are likely responding to the same extrinsic cytokine signal (Fig. 5E). Additionally, these findings demonstrate that the C3 phenotype is not specific to beta cells.

### Visualization of cluster 3-like beta cells in islets from individuals with T1D

To validate these findings in tissues, we performed immunofluorescence on formalin fixed, paraffin embedded (FFPE) pancreatic tissue sections obtained from the Network for Pancreatic Organ Donors with Diabetes (nPOD). We examined 4 T1D donors and 1 ND donor from nPOD, as well as 1 internal ND control (clinical information for these samples is in Supplementary Table 21). We jointly visualized IRF1 and insulin (INS) to identify C3 beta cells (INS+IRF1+) and non-C3 beta cells (INS+IRF1-) that we had found in the above studies. We labeled immune cells with CD45.

We identified a subset of beta cells containing IRF1 in 3 out of 4 T1D donors (Fig. 6, Supplementary Fig. 14A). The T1D sample containing no C3 beta cells (6551) had very few beta cells and insulin-containing islets overall (6579 similarly had few insulin-containing islets).

**Fig. 6:**
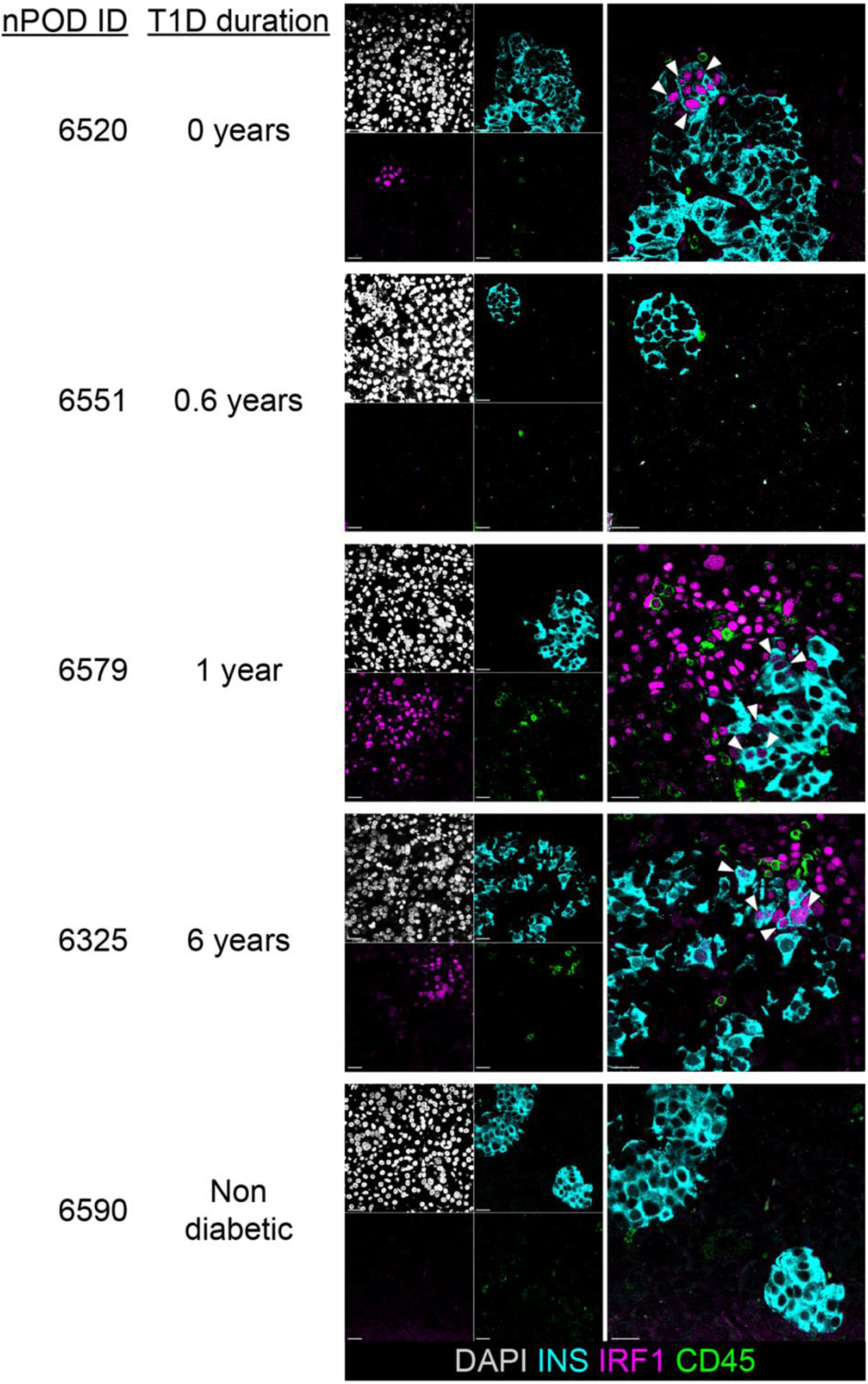
Visualization of C3 beta cells in islets from donors with T1D. INS (cyan), IRF1 (magenta), CD45 (green), and a nuclear marker (DAPI; grey) jointly visualized in FFPE pancreatic tissue sections. Single channel images (left) and composite images (right) are shown. Brightness and contrast were adjusted to enhance visualization. Arrows indicate C3 beta cells (INS+IRF1+ cells). Scale bars correspond to 20μm. Additional clinical information is present in Supplementary Table 21.

Interestingly, within individual islets, C3 beta cells were often adjacent to other C3 beta cells. This spatial distribution is consistent with the local diffusion of secreted cytokines. In the donors with abundant insulin-containing islets (6520 and 6325), despite certain islets have several C3 beta cells, we observed other insulin-containing islets lacking any C3 beta cells, suggesting that these cells are spatially restricted to certain islets within individual donors (Supplementary Fig. 14B). We did not observe any C3 beta cells in either ND sample, aligning with a previous study by Colli et al. which reported absent IRF1 in ND islets (Fig. 6, Supplementary Fig. 14C) (Colli et al., 2018).

## Discussion

Here, we sought to identify the characteristics of beta cells persisting despite ongoing autoimmunity in T1D. We used scRNAseq data from human islets that were generated by HPAP, as well as T1D human tissues from nPOD. We applied a GRN inference-based clustering approach to study subpopulations of surviving beta cells in T1D. While this method is limited by incomplete known TFs and cis-regulatory information, it uncovered a novel population of beta cells, C3, which was enriched in T1D islets. This study highlights the importance of single cell approaches which can identify rare populations often missed by bulk approaches. We found that C3 beta cells expressed genes associated with cytokine responsiveness and immunomodulation, including *SOCS1*, *SOCS3*, and *HLA-E*, and had reduced expression of genes linked to beta cell identity or function, such as the proteases *PCSK1* and *CPE.* Importantly, our findings did not appear to be driven by samples acquired at the time of diagnosis, as we observed a similar or even increased frequency of C3 beta cells in donors with longer disease durations. This transcriptional program was also present in alpha cells that are not eliminated in the autoimmune attack, suggesting it is not limited to beta cells. We confirmed our scRNAseq analyses using patient tissue sections and observed that not only were IRF1+ beta cells present in T1D donors, but these cells were also spatially aggregated. This suggests that the transcriptional profile of C3 beta cells may represent a coordinated response to an inflammatory context.

Several studies have manipulated the genes enriched in C3 and identified roles in beta cell survival. IRF1 is particularly intriguing, as islets lacking *Irf1* were more rapidly eliminated in a murine allograft model (Gysemans et al., 2009). Furthermore, beta cell *IRF1* silencing has been shown to increase production of inflammatory chemokines in several preclinical rodent models (Baker et al., 2003; Moore et al., 2011). Expression of *SOCS1*, which is a known target of IRF1 (Saito et al., 2000), has been identified as a protective factor from cytokine-induced apoptosis in a CRISPR screen of EndoC-BH1 cells (Benaglio et al., 2022). IRF1 has also been shown to induce *CD274* (encoding PDL1), *SOCS3*, and *HLA-E* expression in EndoC-BH1 cells in response to IFNA (Colli et al., 2020). In conjunction with our findings that long-lived T1D beta cells have increased IRF1, it is tempting to speculate that IRF1 may be a critical factor governing beta cell survival in T1D. Intriguingly, *IRF1, SOCS1,* and other genes enriched in C3 (such as *IKZF4* and *BCL6*) have known T1D-associated variants, raising the possibility that this program may be altered in T1D (Benaglio et al., 2022; Onengut-Gumuscu et al., 2025; Robertson et al., 2021).

Other transcription factors predicted to be more active in C3 may likewise influence beta cell survival. CEBPD and JUNB have both been found to protect islets from cytokine-induced apoptosis in vitro (Gurzov et al., 2012; Gurzov et al., 2008; Moore et al., 2012). BCL6 has also been shown to be cytokine-responsive in human beta cells (Benaglio et al., 2022).

Overexpression of human BCL6 in beta-like INS-1E cells was found to increase apoptosis while decreasing the production of inflammatory factors after cytokine stimulation (Igoillo-Esteve et al., 2011). Finally, ETV6 represses expression of inflammatory genes in human peripheral blood mononuclear cells (Fisher et al., 2020). Altogether, the co-expression of these transcription factors is suggestive of an immunomodulatory program which is active in residual T1D beta cells.

We also observed decreased expression of certain functional proteins in both C3 beta cells and C3 alpha cells, such as *SLC30A8, PCSK1/2* (depending on the cell type), and *SCGN.* Many of these genes encode autoantigens in beta cells, which may influence the immunogenicity of C3 beta cells. It is also possible that increased frequencies of C3 beta cells and/or C3 alpha cells may explain known secretory defects in T1D patients (Brissova et al., 2018; Huber et al., 2025; Krogvold et al., 2015; O’Meara et al., 1995). For instance, *PCSK1/2* mediate proinsulin cleavage, and an elevated serum proinsulin:insulin ratio is a hallmark of T1D. Insulin secretion has been shown to improve in T1D islets after several days of culture ex vivo (Krogvold et al., 2015), perhaps due to the resolution of cytokine-induced secretory impairment in C3-like beta cells.

We propose that this cluster (C3) represents a resilient group of cells akin to the beta cell population (“Btm” cells) we previously identified as resistant to killing in NOD mice (Rui et al., 2017). Several similarities exist between C3 beta cells and Btm cells, including enrichment in autoimmune diabetes, loss of cellular identity, and evidence of cytokine signaling. Cytokine responsiveness in immune resilient Btm cells was a paradoxical finding given that cytokines can be cytotoxic to beta cells (Eizirik et al., 2021). However, adaptive cytokine effects have recently been documented in vitro, where priming with low-dose IL1B protected INS-1E cells from death upon exposure to higher dose IL1B and IFNG (Sétula et al., 2025). In light of these findings, it is possible that sublethal cytokine sensing drives the C3 state to promote beta cell survival in human T1D patients. We emphasize that this is one possible interpretation which cannot be validated solely from cross-sectional analyses, which we performed here. An alternative explanation is that the enrichment of C3 in T1D donors is associative, reflecting a response to the inflammatory milieu present in T1D islets rather than C3 causally driving beta cell survival.

While the identification of this phenotype among surviving beta cells is an important advance, future studies directly testing if the C3 state alters beta cell death or survival are required to clarify the functional relevance of these cells.

There are several limitations to our study. The fate of the AAb+ individuals is unknown and therefore our findings in the islets from these individuals may reflect progression or a protective mechanism if the individuals did not progress. In addition, by definition all of the T1D beta cells that we identified are survivors. We do not know the phenotype of cells that were destroyed during progression, nor do we know the cell death mechanisms that surviving cells may have endured. We also do not know which cells produce the cytokines likely driving beta cells to adopt the C3 state, making this an important future direction. Finally, our inference-based approach did not directly test the activity of specific factors compared to other methods (e.g. ChIP-seq).

In summary, we identified a novel population of beta cells in T1D that likely arises due to sensing of inflammatory cytokines. To our knowledge, this is the first time a subtype of beta cells has been identified in T1D using unbiased methods. We hypothesize that the transcriptional network active in these cells may influence beta cell survival and inform mechanisms of T1D pathogenesis. If the C3 phenotype indeed confers resistance to immunological attack, our findings identify key targets for engineering more durable beta cells for the treatment of patients with established T1D.

## Author contributions

MS designed, conducted studies, analyzed the data and wrote the manuscript. ZT and JSP performing tissue staining and image acquisition and wrote the manuscript. JST and KCH designed studies, reviewed data, and wrote the manuscript.

## Acknowledgements

The authors would like to thank the donors and their families for their generous contributions. We also thank the members of the HPAP for their considerable efforts to generate the dataset used in this study. We thank the Yale Center for Research Computing for their support and guidance.

## Declaration of Competing Interests

J.S.T. serves on the scientific advisory boards of CytoReason Inc. and Immunoscape Inc. and as the co–chief scientific officer (as an unpaid volunteer) of the Human Immunome Project. K.C.H. has a patent for the use of teplizumab for delay of type 1 diabetes (patent US20220041720, issued February 10, 2022) but no financial interest.

## Funding support

This work is the result of NIH funding, in whole or in part, and is subject to the NIH Public Access Policy. Through acceptance of this federal funding, the NIH has been given a right to make the work publicly available in PubMed Central. Support from T32GM136651, and DK057846, DK129523 from the NIH and 2-SRA-2022-1197-S-B from Breakthrough T1D, to KCH. In addition JST has received a CZ Biohub NY Investigator Award.

## Data Availability

All data analyzed in this study is publicly available. HPAP scRNAseq data can be downloaded from https://hpap.pmacs.upenn.edu/. Maestas et al. in vitro islet scRNAseq and snMultiome data can be downloaded from GSE237448. Benaglio et al. in vitro beta cell snATACseq data (peaks aggregated by sample) can be downloaded from GSE205853. The file“GSE205853_beta.snATAC_count.matrix.txt.gz” was used in this study. Code and Supplementary Tables 1-20 are available at: https://doi.org/10.5281/zenodo.19934213.

## Methods

### HPAP scRNAseq data generation

The generation of scRNAseq data by HPAP has been described previously (Kaestner et al., 2019). Briefly, islets were isolated then cultured, handpicked, and dissociated into a single cell suspension. A detailed methodology for this process is available at https://hpap.pmacs.upenn.edu/explore/workflow/islet-molecular-phenotyping-studies?protocol=4. Library preparations were then generated using 10X Genomics Chromium Single Cell 3’ Reagent Kit (either version v2, v3, or v3.1 depending on the donor). Fastq files from 56 donors who were either T1D (n = 12), AAb+ (n = 11) or ND (n = 33) were downloaded directly from PANC-DB (https://hpap.pmacs.upenn.edu/explore/download?matrix). Count matrices were generated using the CellRanger count function from 10X Genomics CellRanger version 9.0.1, aligning to the GRCh38 2020-A reference genome available from https://www.10xgenomics.com/support/software/cell-ranger/downloads#reference-downloads.

Sample and assay metadata is found in Supplementary Table 1 (reference data was downloaded from https://hpap.pmacs.upenn.edu/). T1D donors HPAP-064, HPAP-084, and HPAP-113 did not have annotated disease durations on PANC-DB. We confirmed that these were recent-onset T1D cases by contacting HPAP.

### HPAP scRNAseq processing

Analysis was performed in R (version 4.4.1) with Seurat (version 5.1.0) (Hao et al., 2024). Cells were initially filtered for those with between 500 and 10000 unique genes detected and fewer than 15% mitochondrial transcripts. ND donor HPAP-093 was removed due to disproportionately low median UMI per cell (as in Elgamal et al.) (Elgamal et al., 2023). SoupX (version 1.6.2) was used for correction of ambient transcripts using the automated method for estimating contamination fraction and rounding of corrected counts to integers (Young & Behjati, 2020). The default CellRanger clustering labels for each sample were provided to SoupX. scDblFinder (version 1.18.0) was used to identify and remove likely doublets on a per sample basis with default parameters (Germain et al., 2021).

### HPAP scRNAseq islet clustering

Data was normalized using the SCTransform v2 method with regression of percent of mitochondrial transcripts (Choudhary & Satija, 2022). Principal component (PC) analysis was performed using RunPCA, then data was integrated across samples using the canonical correlation analysis (CCA) method (Butler et al., 2018). The first 30 CCA dimensions were used for Seurat FindNeighbors and RunUMAP, and FindClusters was run at a resolution of 0.15.

Marker genes for each cluster were identified using PrepSCTFindMarkers and FindAllMarkers (default parameters, Wilcoxon Rank Sum test, Supplementary Table 2). A beta cell cluster was defined by the presence of known marker genes, such as *INS* and *IAPP*. Three alpha cell clusters were also identified. Both cell types were isolated and processed downstream, independently by cell type.

### Post-islet clustering refinement of beta and alpha cells

Each cell type was renormalized using the SCTransform v2 method with regression of percent of mitochondrial transcripts. Doublets were then manually removed from the beta cell cluster by removing cells highly expressing known marker genes for other pancreatic cell type (alpha: *GCG*, delta: *SST*, gamma: *PPY*, epsilon: *GHRL*, acinar: *REG1A*, endothelial: *PLVAP*, ductal: *CFTR*, stellate: *PDGFRB*, T: *CD3D*, myeloid: *C1QC*, or mast: *KIT*) (Melton et al., 2025). Alpha cells were filtered using the same thresholds (removing *INS* expressing cells instead of *GCG* expressing cells). UCell (version 2.8.0) scores were calculated for remaining cells using gene sets contained in Supplementary Table 3 (Andreatta & Carmona, 2021). Gene sets were curated from Hallmark, Gene Ontology (GO) Biological Process, Reactome, KEGG, and several individual publications (Aleksander et al., 2023; Ashburner et al., 2000; Cheung et al., 2023; Jassal et al., 2020; Kanehisa & Goto, 2000; Liberzon et al., 2015; McKinney et al., 2015; Subramanian et al., 2005; Wherry et al., 2007; Woods et al., 2013). Gene sets were filtered to those with at least 10 and no more than 300 genes. UCell scores were appended Seurat objects. Clustering on UCell scores was performed for both beta and alpha cells using the first 15 calculated PCs (after RunPCA) for FindNeighbors and RunUMAP, and FindClusters was run at a resolution of 0.15. A population of cells with absent *INS* or *GCG* (for beta and alpha cells respectively) and elevated exocrine markers was discarded (Supplementary Fig. 3A-G for beta cells and Supplementary Fig. 12 for alpha cells). Additional filtering was applied after GRN inference to produce the final populations of beta and alpha cells used for all HPAP cell analyses (see section “Clustering on regulon scores”). Note that for the cell type frequency plots in Supplementary Fig. 2B&C, the post-filtering population was used for beta cells while the pre-filtering population was used for alpha cells.

### GRN inference

GRN inference was performed using the Python (version 3.10.0) implementation of SCENIC (pySCENIC, version 0.12.1) on beta and alpha cells independently (Aibar et al., 2017; Van de Sande et al., 2020). Natural log + 1 transformed SoupX-corrected counts were used as input. Reference files were downloaded from the cisTarget repository (https://resources.aertslab.org/cistarget/). The most recent GRCh38 databases and motif annotations (v10) were used. Gene filtering was performed to select genes detected in at least 5 cells and genes with symbols overlapping in the reference SCENIC databases. The adjacency matrix for pySCENIC was constructed using RegDiffusion (version 0.1.1) (Zhu & Slonim, 2024). Edges in the top 50% by weight were extracted. For each gene, the top 20 edges were considered. Regulons were pruned using pySCENIC ctx (complete output of this step for beta cells is in Supplementary Table 5). Regulon activity scores were calculated using AUCell with default parameters. This was repeated 9 additional times for beta cells (along with cell filtering described at the start of the next section) to create the pairwise correlation matrices in Supplementary Fig. 7. AUCell regulon scores were then exported and appended to the beta or alpha cell Seurat objects for downstream use. For beta cells, the AUCell scores from the initial model training were used.

### Clustering on regulon scores

After pySCENIC GRN inference, cells with less than 1000 unique genes were removed. Remaining beta cells comprised the finalized beta cell object. For clustering, regulon scores were rank normalized then scaled across cells to account for certain regulons being zero inflated and/or having long tails. RunPCA was performed on transformed regulon scores (beta cell PC4 scores in Fig. 2I calculated in this step). For beta cells, the first 10 PCs were used for RunUMAP and FindNeighbors based on estimation from an elbow plot, and FindClusters was performed at a resolution of 0.3. The first 8 PCs minus PC6 were used in alpha cells for RunUMAP and FindNeighbors based on estimation from an elbow plot, and FindClusters was performed at a resolution of 0.25. PC6 was excluded due to its correlation with feature count. Differential regulon activity between cell clusters was calculated with Seurat FindAllMarkers using non-rank normalized AUCell scores with mean.fxn set to rowMeans and min.pct and logfc.threshold both set to 0 (Supplementary Table 7 for beta cells and Supplementary Table 17 for alpha cells).

PrepSCTFindMarkers was run on the full post-islet clustering cell objects (i.e. where SCTransform was rerun for beta and alpha cells before cell filtering) prior to differential gene expression testing between cell clusters. Differential gene expression testing between beta and alpha cell clusters was calculated using Seurat FindAllMarkers with min.pct and logfc.threshold set to zero and recorrect_umi set to false (Wilcoxon Rank Sum Test, Supplementary Table 8 for beta cells and Supplementary Table 18 for alpha cells). To specifically identify positive and negative DEGs in C3 for beta and alpha cells, each population was tested against other beta or alpha cells not belonging to that cluster using Seurat FindMarkers with min.pct and logfc.threshold set to zero and recorrect_umi set to false (Wilcoxon Rank Sum Test, Supplementary Table 9 for beta cells and Supplementary Table 19 for alpha cells).

### Gene set enrichment analysis

Gene set enrichment analysis (GSEA) was performed with the fgsea package (version 1.30.0) using gene sets contained in Supplementary Table 3 (described in above “Post-islet clustering refinement of beta and alpha cells” section) (Korotkevich et al., 2021; Mootha et al., 2003; Subramanian et al., 2005). Beta cell C3 DEGs (Supplementary Table 9) or alpha cell C3 DEGs (Supplementary Table 19) were independently filtered to those expressed by at least 1% of cells in one or both test groups (e.g. 1% of cells in C3 or non-C3 beta cells) and ranked by average log2 fold change. The full GSEA outputs are located in Supplementary Table 10 (beta cells) and Supplementary Table 20 (alpha cells).

### Differential RNA expression testing by clinical status

Clinical statuses were compared using a pseudobulk approach on the finalized beta cell object (see above section “Clustering on regulon scores”). SoupX-corrected RNA counts were summated by donor (either for all beta cells or within a single beta cell cluster). A minimum of 5 beta cells was required for donor inclusion in a given comparison. Pairwise differential activity testing between T1D, AAb+, and ND was performed using edgeR (Robinson et al., 2010). Lowly expressed genes were removed from a given comparison using filterByExpr. glmQLFit was used to fit the model ∼ 0 + Clinical Status + Donor Age + Donor Sex + Donor BMI + 10X Chromium Kit Version. Contrasts were calculated using glmQLFTest. Results from all comparisons were merged (Supplementary Table 4). Local and global FDRs (within a contrast and across all contrasts, respectively) were calculated using the BH procedure. Local contrast FDR values were reported in the main text.

### Differential regulon activity testing by clinical status

Clinical statuses were compared using a pseudobulk-like approach on the finalized beta cell object (see above section “Clustering on regulon scores”). Non-rank normalized AUCell scores were averaged by donor (either for all beta cells or within a single beta cell cluster). A minimum of 5 beta cells was required for donor inclusion in a given comparison. Pairwise differential activity testing between T1D, AAb+, and ND was performed using lm and a model of ∼ 0 + Clinical Status + Donor Age + Donor Sex + Donor BMI + 10X Chromium Kit Version.

Contrasts were calculated using the emmeans function from the emmeans package (version 2.0.1). Results from all comparisons were merged (Supplementary Table 6). Local and global FDRs (within a contrast and across all contrasts, respectively) were calculated using the BH procedure. Local contrast FDR values were reported in the main text.

### scRNAseq reanalysis of Maestas et al. in vitro treated primary human islets

Data was downloaded from GSE237448. Primary islets from n = 5 ND human donors were stimulated with the following conditions: IFNG, TNFA, IL1B, IFNG+IL1B, IFNG+IL1B+TNFA (cytokine mix, CM), Brefeldin A, Thapsigargin, or DMSO (unstimulated control) in vitro for 48 hours as described previously (Maestas et al., 2024). Samples from donors 1, 2, and 3 were labeled with hashing antibodies and processed together (within each donor). Samples from donors 4 and 5 were fixed and processed separately. Initial quality control filtering of cells was repeated using the thresholds from the original publication: 2000 ≤ unique genes ≤ 7500 (donor 1); 1000 ≤ unique genes ≤ 7500 (donor 2); 1000 ≤ unique genes ≤ 6000 (donor 3); 200 ≤ unique genes ≤ 9000 (donors 4&5). For hashed samples, hashed oligos were normalized using a CLR transformation and demultiplexed using HTODemux. Singlets were selected for downstream use. All samples were then merged and normalized using the SCTransform v2 method with regression of percent of mitochondrial transcripts. Samples were integrated using Harmony (version 1.2.0) after RunPCA (Korsunsky et al., 2019). Clustering of islet cells was performed using the first 30 Harmony dimensions for FindNeighbors and RunUMAP, and FindClusters was run at a resolution of 0.3. DEGs for each islet cell cluster were calculated with PrepSCTFindMarkers and FindAllMarkers (Wilcoxon Rank Sum Test, Supplementary Table 11). Clusters M1, M4, M7, M8, and M15 expressed beta cell markers (e.g. *INS*, *IAPP*, *MAFA*) were defined as beta cells (Supplementary Fig. 10D-E and Supplementary Table 11). Cluster M12 expressed *INS* but was composed of low-quality cells (low unique gene abundance) and was thus not included (Supplementary Fig. 10F).

### Differential expression testing in Maestas et al. scRNAseq in vitro beta cells

DEGs were calculated using edgeR (Robinson et al., 2010). Transcript counts were summated within each sample (combination of donor and treatment condition). Lowly expressed genes within a treatment condition were removed with filterByExpr. glmQLFit was used to fit the model ∼ 0 + Treatment Condition + Donor. Contrasts were calculated using glmQLFTest.

Results from all comparisons were merged (Supplementary Table 13). Local and global FDRs (within a contrast and across all contrasts, respectively) were calculated using the BH procedure. Local contrast FDR values were reported in the main text.

### Scoring cluster 3 gene sets in Maestas et al. scRNAseq in vitro beta cells

Transcript counts were summated within each sample (combination of donor and treatment condition). Lowly expressed genes within a treatment condition were removed with edgeR filterByExpr. Aggregated counts were normalized with log2(counts per million + 1). Gene sets representing cluster 3 up and cluster 3 down genes were created by taking the top 50 non-ribosomal/mitochondrial genes by adjusted p value with log fold changes above and below 0, respectively (from Supplementary Table 9). Aggregated normalized counts were scored for activity of these gene sets using GSVA (version 1.52.3) with kcdf set to Gaussian (Hänzelmann et al., 2013). Differential GSVA testing was performed using limma (version 3.60.3) lmfit with the model ∼ 0 + Treatment Condition + Donor (Ritchie et al., 2015). Contrasts were calculated using eBayes with contrasts.fit. Results from all treatment conditions were merged (Supplementary Table 12). A global FDR was calculated from nominal p values across all comparisons with the BH procedure and was used in the main text.

### snMultiome reanalysis of Maestas et al. in vitro treated primary human islets

Data was downloaded from GSE237448. Primary islets from n = 1 ND human donor were treated with IFNG, TNFA, and IL1B (CM) or control using the same duration and concentrations as for the scRNAseq experiments. Signac (version 1.13.0) was used for analysis (Stuart et al., 2021). Initial quality control filtering of cells was repeated using the thresholds from the original publication: 1000 ≤ RNA transcripts ≤ 50000, 1000 ≤ ATAC fragments ≤ 60000, nucleosome signal < 0.15 and transcription start site (TSS) enrichment > 0.2. Peak calling was done with MACS2 (version 2.2.9.1) using hg38 reference and EnsDb.Hsapiens.v86 annotation. RNA data was processed using SCTransform and Harmony (as for Maestas et al. scRNAseq data). ATAC data was processed using FindTopFeatures, RunTFIDF and RunSVD. Integrated clustering and visualization was performed using FindMultiModalNeighbors (with the first 50 Harmony dimensions and 2-40^th^ ATAC dimensions) and FindClusters with a resolution of 0.3. Cluster DEGs in Supplementary Table 14 were calculated with PrepSCTFindMarkers and FindAllMarkers. To score motif accessibility, chromVAR was run using the core JASPAR2020 motif database on peaks mapping to chromosomes (i.e. not scaffolds) (Fornes et al., 2020; Schep et al., 2017). Beta cells were identified by cells expressing known beta cell marker genes (Supplementary Fig. 11; cluster A6 was excluded due to low RNA quality). Differential motif accessibility between CM and CTRL treated beta cells was performed using chromVAR z-scores and FindMarkers with mean.fxn set to rowMeans, min.pct set to 0 and logfc.threshold set to 0 (Wilcoxon Rank Sum Test, Supplementary Table 15).

### Identification and motif analysis of differentially accessible snATAC peaks

Experimental data was generated in Benaglio et al. (Benaglio et al., 2022). Primary islets from non-diabetic human donors (n = 3) were either treated with 10 ng/mL IFNG, 0.5 ng/mL IL1B and 1 ng/mL TNFA or unstimulated for 24 hours in vitro. snATACseq was performed using the 10X Genomics Chromium Single Cell ATAC kit. Peak calling and clustering were performed to resolve islet cell populations, and aggregated count matrices were generated for each sample (donor and treatment condition pair) by cell type as previously described. Aggregated beta cell peak counts were downloaded from GSE205853. As in Benaglio et al., differentially accessible peaks between stimulated versus unstimulated beta cells were identified using DESeq2 (version 1.44.0) with a model of ∼ Treatment Condition + Donor (Love et al., 2014). Differentially accessible peaks were selected by those with a BH-calculated FDR < 0.10 (as in the original publication). Motif analysis of differentially accessible peaks was done using HOMER (version 5.1) and the hg19 human genome assembly (hg19 was used when generating the count matrix in Benaglio et al.) (Heinz et al., 2010). De novo motifs were identified in differentially accessible peaks using findMotifsGenome.pl with the size argument set to given (full table of identified TF motifs is found in Supplementary Table 16).

### Immunofluorescence and image acquisition

Formalin-fixed, paraffin-embedded (FFPE) pancreatic tissue sections were obtained from nPOD or the Yale Pathology Tissue Service. FFPE sections were deparaffinized and antigen retrieval was performed using a Tris-based antigen unmasking solution (Vector Laboratories, #H-3301-250). Immunostaining was performed with rabbit anti-IRF1 (Cell Signaling Technology, #8478, 1:100 dilution), mouse anti-CD45 (Lunaphore, #MR10090, 1:100 dilution) and guinea pig anti-insulin (Invitrogen, #180067, 1:400 dilution) antibodies. Fluorescently conjugated secondary antibodies were used at a 1:200 dilution (Invitrogen, #A-32790, #A-31570, and #A-21450, respectively). Additionally, DAPI nuclear staining was performed. Imaging was performed using a Stellaris 5 confocal microscope (Leica Microsystems).

### Data visualization

Plots were generated using ggplot2 (version 3.5.1), ggh4x (version 0.2.8), ggrepel (version 0.9.5) and ggsignif (version 0.6.4). The heatmaps in Fig. 3A and Supplementary Fig. 7 were generated using ComplexHeatmap (version 2.20.0). The cluster tree in Supplementary Fig. 5 was generated using clustree (version 0.5.1) (Zappia & Oshlack, 2018).

---

**Supplementary Fig. 1:**
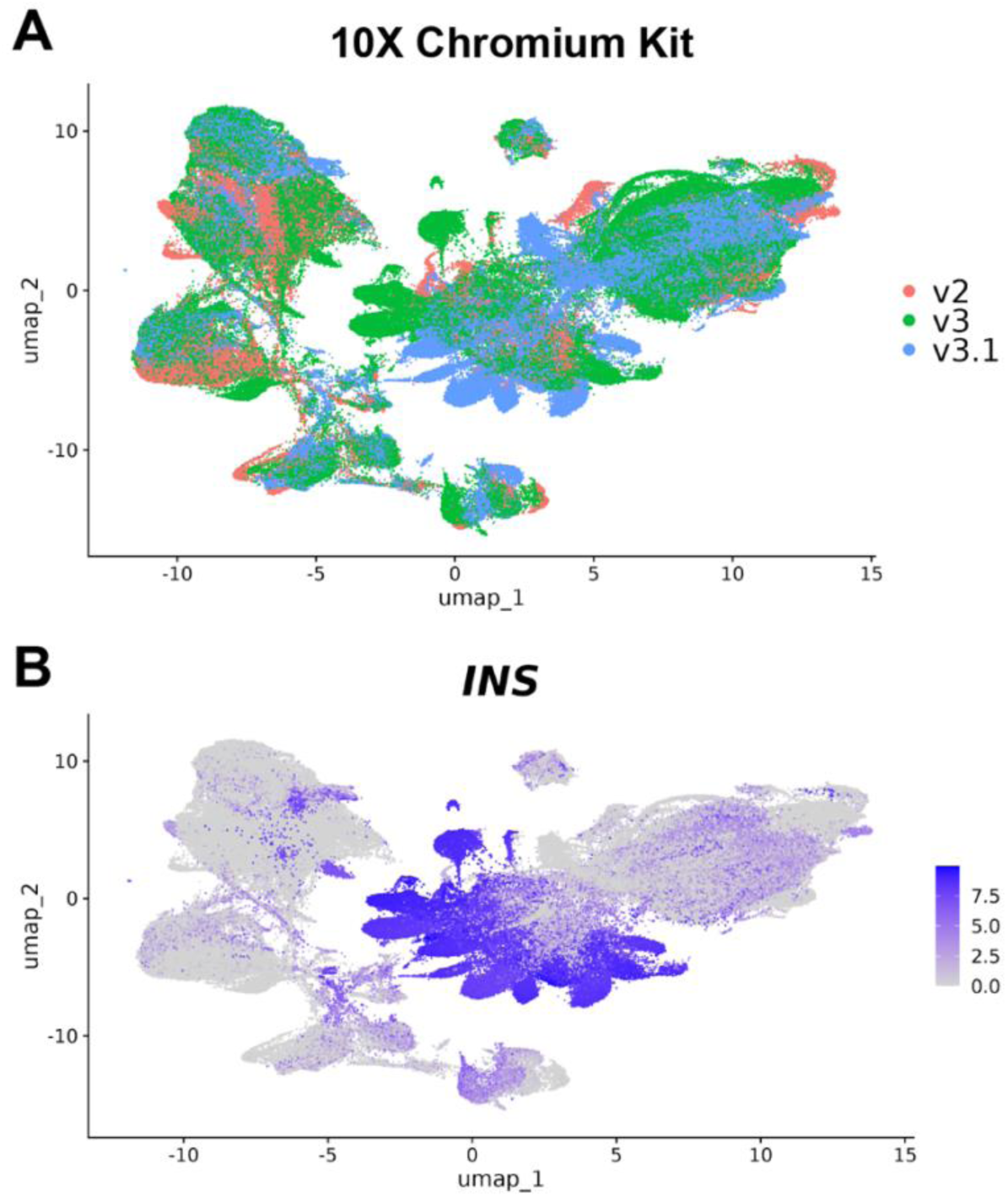
Technical differences in HPAP islet scRNAseq dataset. A: Unintegrated UMAP plot of HPAP islet cells after dimensionality reduction (PCA on normalized transcript counts). Cells are colored by 10X Chromium kit chemistry. B: Feature plot of *INS* expression to highlight beta cells. This unintegrated UMAP was not utilized for any downstream analyses.

**Supplementary Fig. 2:**
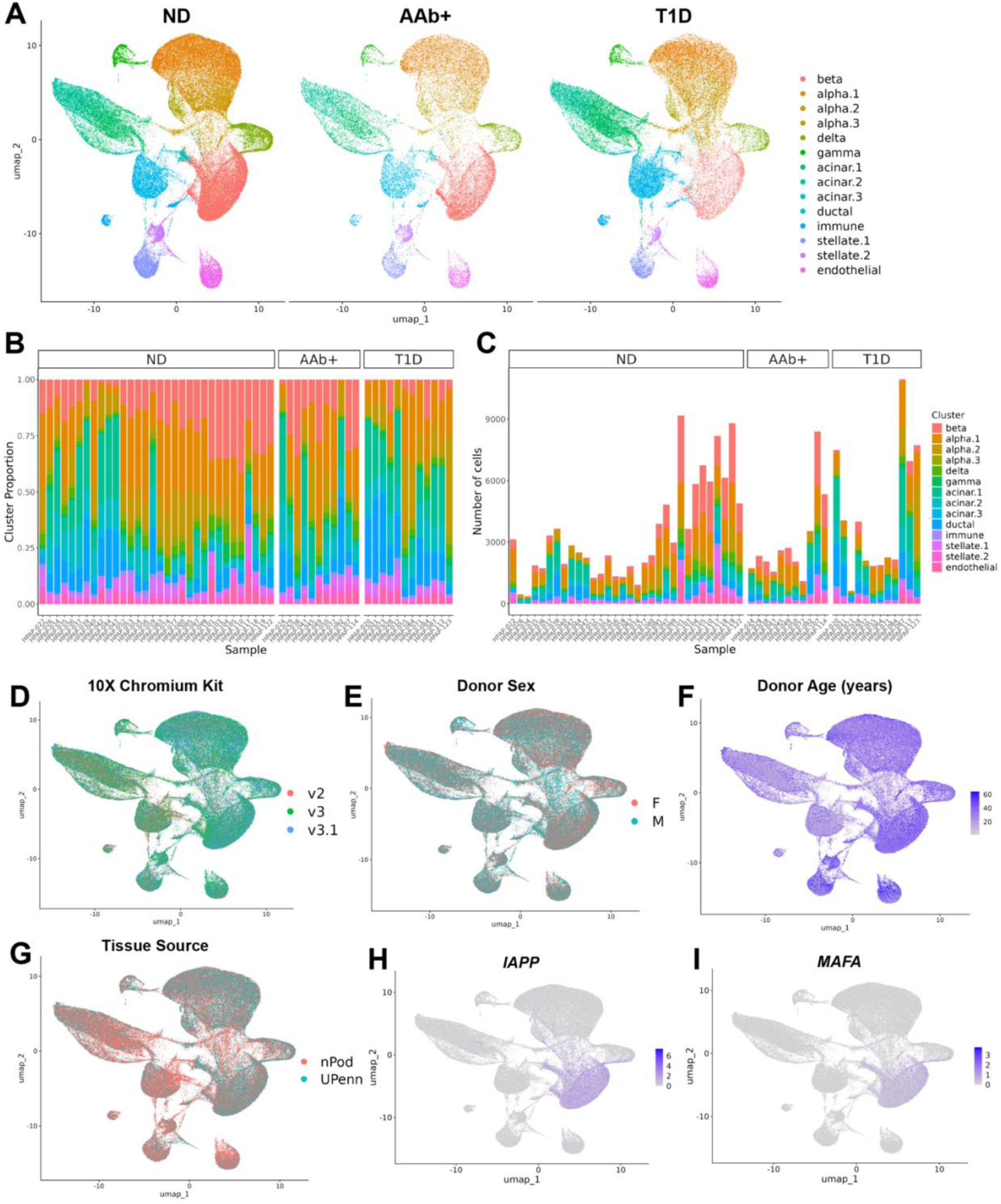
Identification of beta cells in T1D, AAb+, and ND donors from HPAP scRNAseq data. A: Integrated UMAP of HPAP islet cell types split by clinical status. B-C: Stacked bar plot showing frequency (B) and count (C) of each islet cell type per donor. Beta cell frequencies/counts shown here are those following post-islet clustering refinement and low feature count filtering of the beta cell population (Methods). D-G: Integrated UMAP of islet cell types by 10X Chromium reagent kit chemistry (D), donor sex (E), donor age (F), and tissue source (G). H-I: Feature plots of islet expression of *IAPP* (H) and *MAFA* (I).

**Supplementary Fig. 3:**
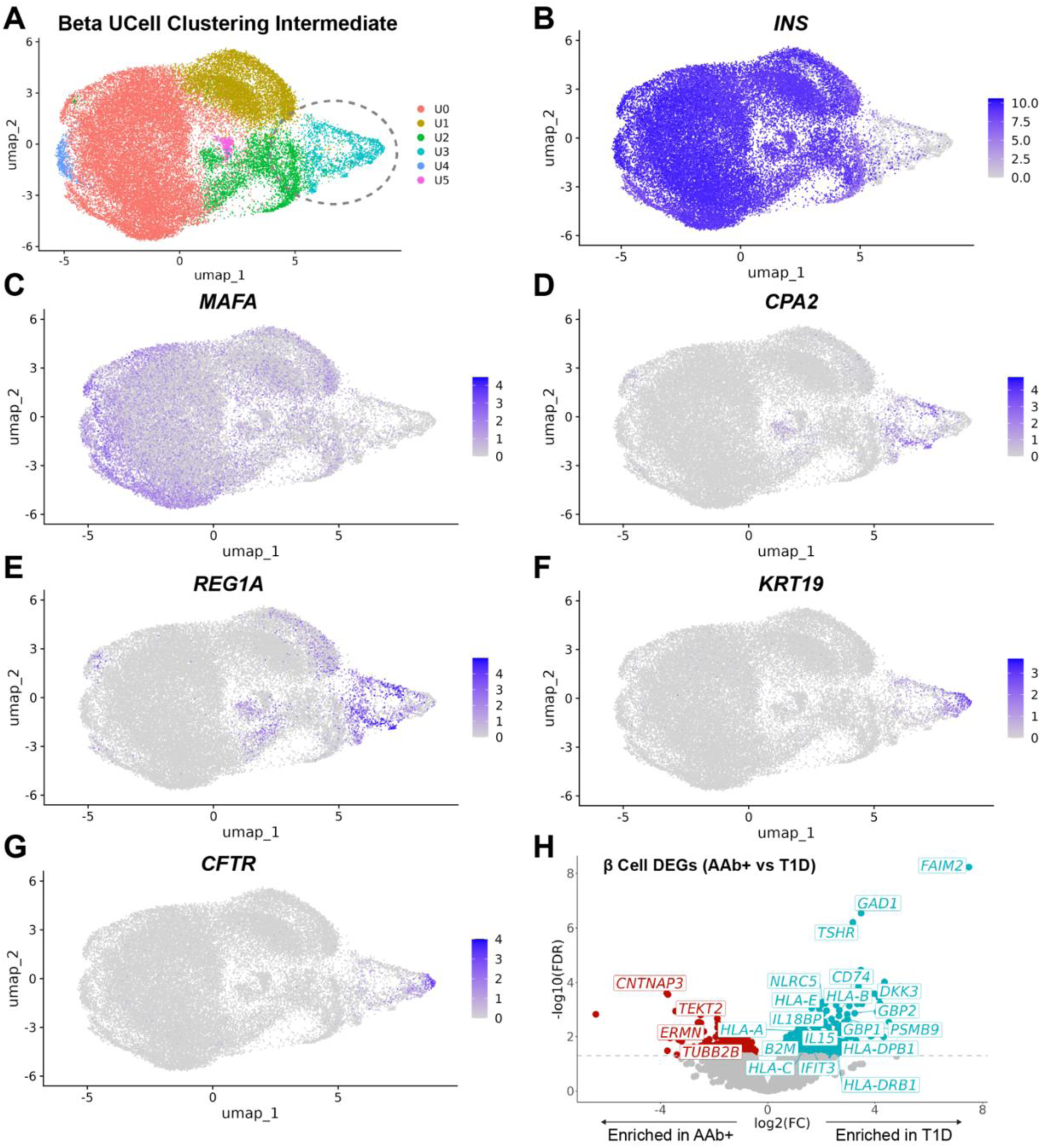
Intermediate beta cell UCell clustering. A: Intermediate beta cell clustering based on UCell scores (Methods). Gene sets for UCell score calculation are in Supplementary Table 3. Cells in cluster U3 (circled) were identified as likely contaminants and removed. B-G: Feature plots showing normalized expression of indicated transcripts. H: Volcano plot of DEGs between T1D and AAb+ beta cells calculated using edgeR with donor age, sex, BMI and 10X Chromium kit chemistry as covariates (as in Fig. 1F). A pseudobulk method was used by aggregating counts in beta cells by donor (Methods). BH-calculated FDR values are shown. The dashed line indicates FDR = 0.05.

**Supplementary Fig. 4:**
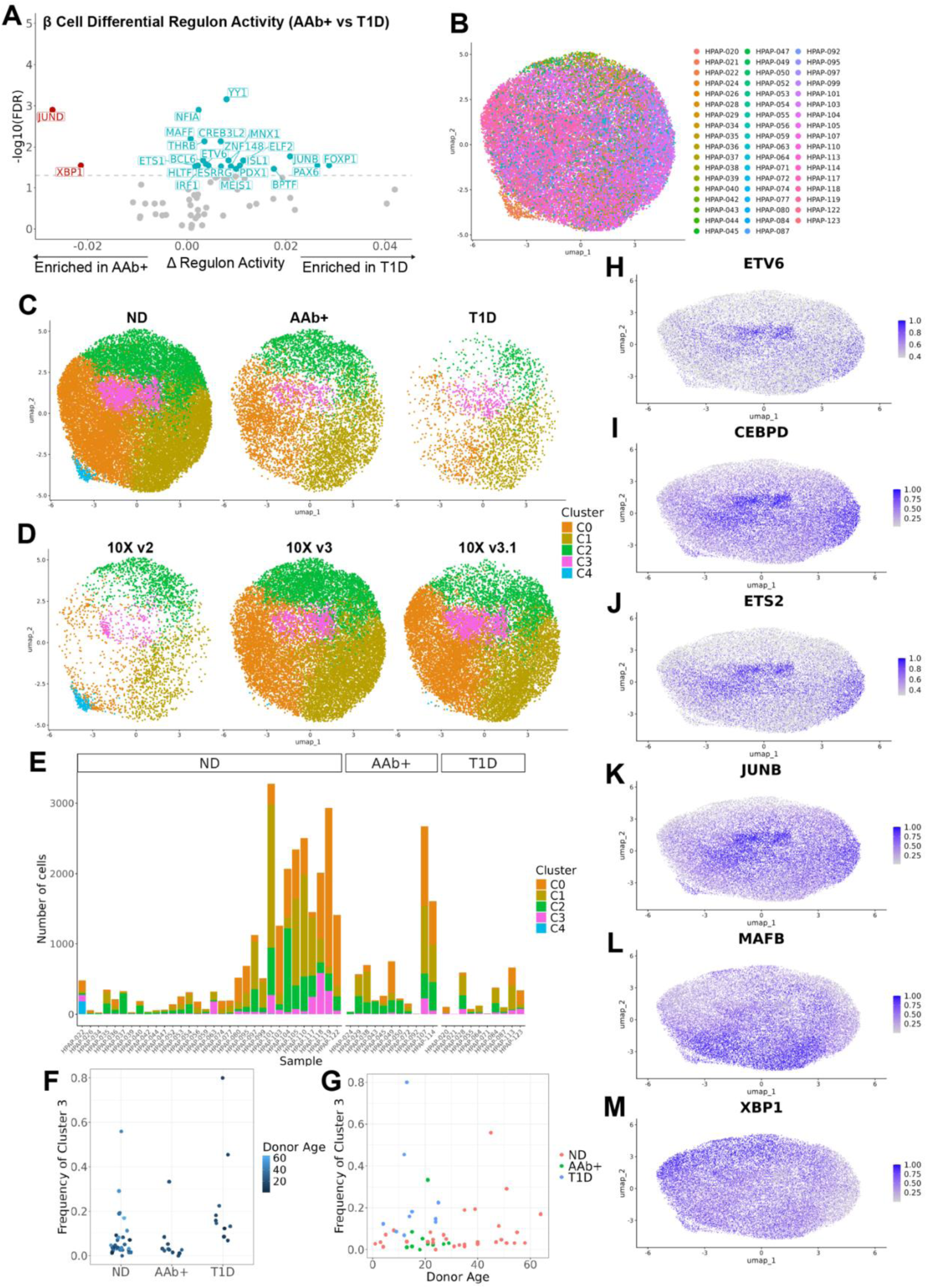
Identification of a T1D enriched beta cell population through GRN inference-based clustering. A: Volcano plot of differentially active regulons between T1D and AAb+ beta cells (as in Fig. 2A). Regulon scores were averaged by donor. Differential activity testing was performed using a linear model with donor age, sex, BMI and 10X Chromium kit chemistry as covariates. BH-calculated FDR values are shown. The dashed line indicates FDR = 0.05. B: Beta cell UMAP by individual donor. C-D: Beta cell UMAP split by clinical status (C) and 10X Chromium reagent kit chemistry (D). E: Count of each beta cell cluster per donor. F: Frequency of C3 beta cells (among all beta cells) by clinical status. Each dot is one donor, shaded by donor age. G: Frequency of C3 beta cells (among all beta cells) versus donor age. Each dot is one donor, colored by clinical status. H-M: Feature plots of rank-normalized regulon activity scores.

**Supplementary Fig. 5:**
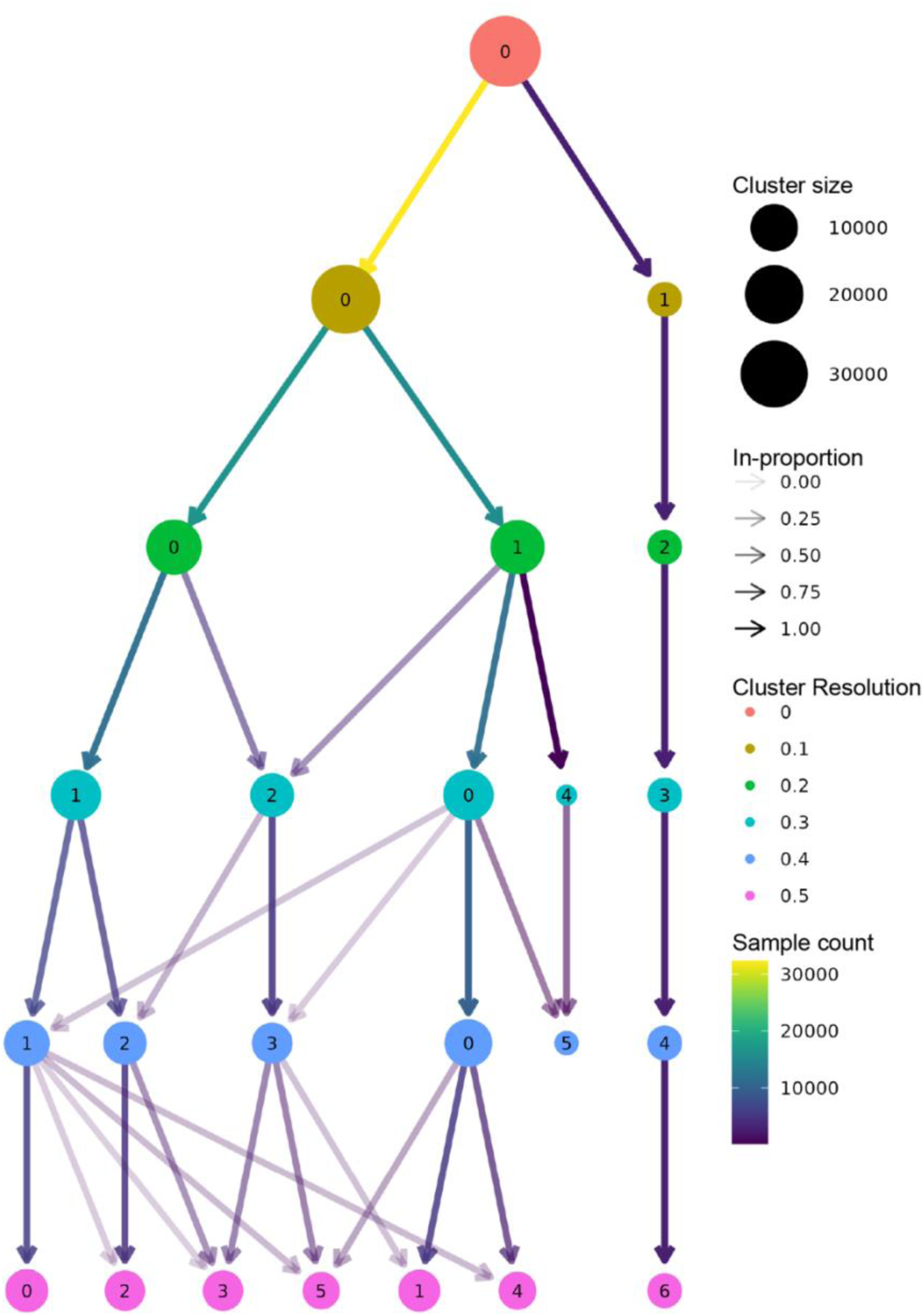
Beta cell cluster 3 is stable across clustering resolutions. Clustree plot showing clustering solutions at different resolutions (Zappia & Oshlack, 2018). A resolution of 0.3 was used for the UMAP in Fig. 2B. Edges from parent nodes contributing less than 10% of cells to the child node are not shown.

**Supplementary Fig. 6:**
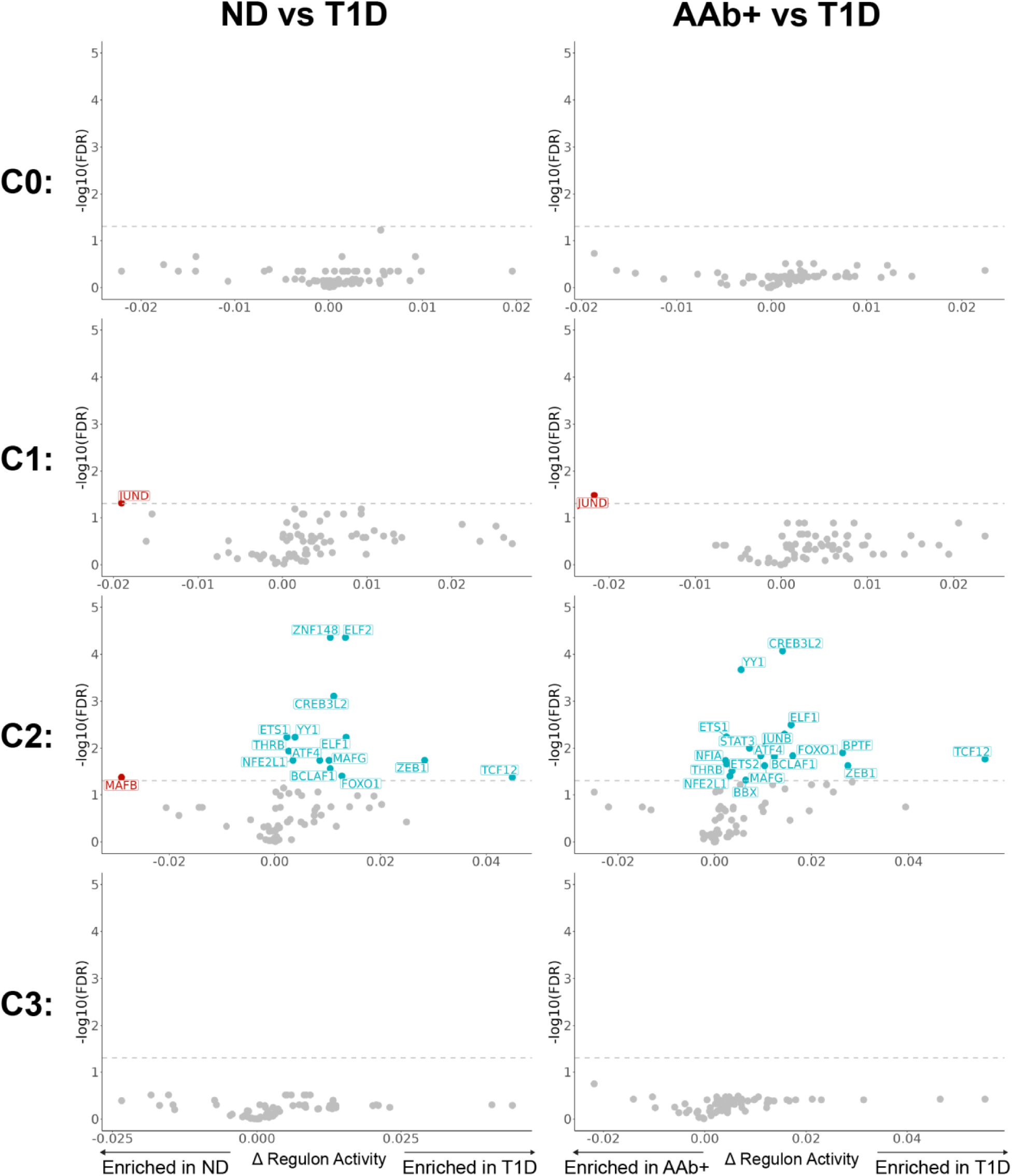
Differential regulon activity between T1D and ND/AAb+ beta cells within each beta cell subcluster. Volcano plots of differentially active regulons between T1D and ND/AAb+ beta cells for each beta cell cluster. Regulon scores were averaged by donor for each cluster. Donors with less than 5 beta cells in a particular cluster were excluded for comparisons within that cluster. Differential activity testing was performed using a linear model with donor age, sex, BMI and 10X Chromium kit chemistry as covariates. BH-calculated FDR values are shown on the y axis. Dashed lines indicate FDR = 0.05.

**Supplementary Fig. 7:**
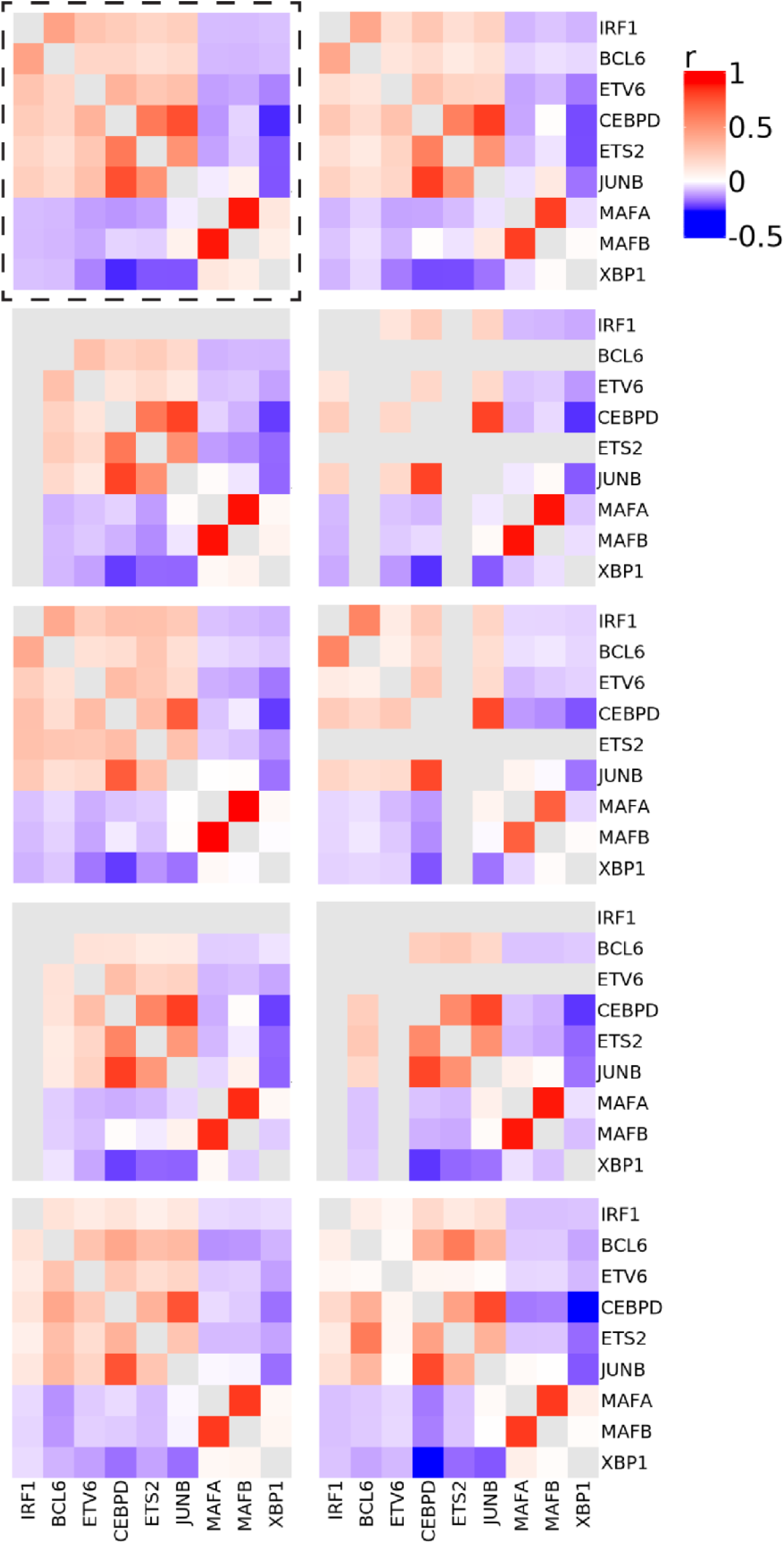
Stability of RegDiffusion model for SCENIC GRN inference. Regulons were scored after independently re-training the RegDiffusion model 9 additional times on the same beta cell population as used for the original run (original run used for all analyses in the dashed box, 10 total trainings). Pairwise Pearson correlation matrices for beta cell activity of the top 6 up and top 3 down C3-enriched regulons are shown. Greyed rows/columns indicate that the regulon was not detected in a given run. The matrix diagonal is also colored grey.

**Supplementary Fig. 8:**
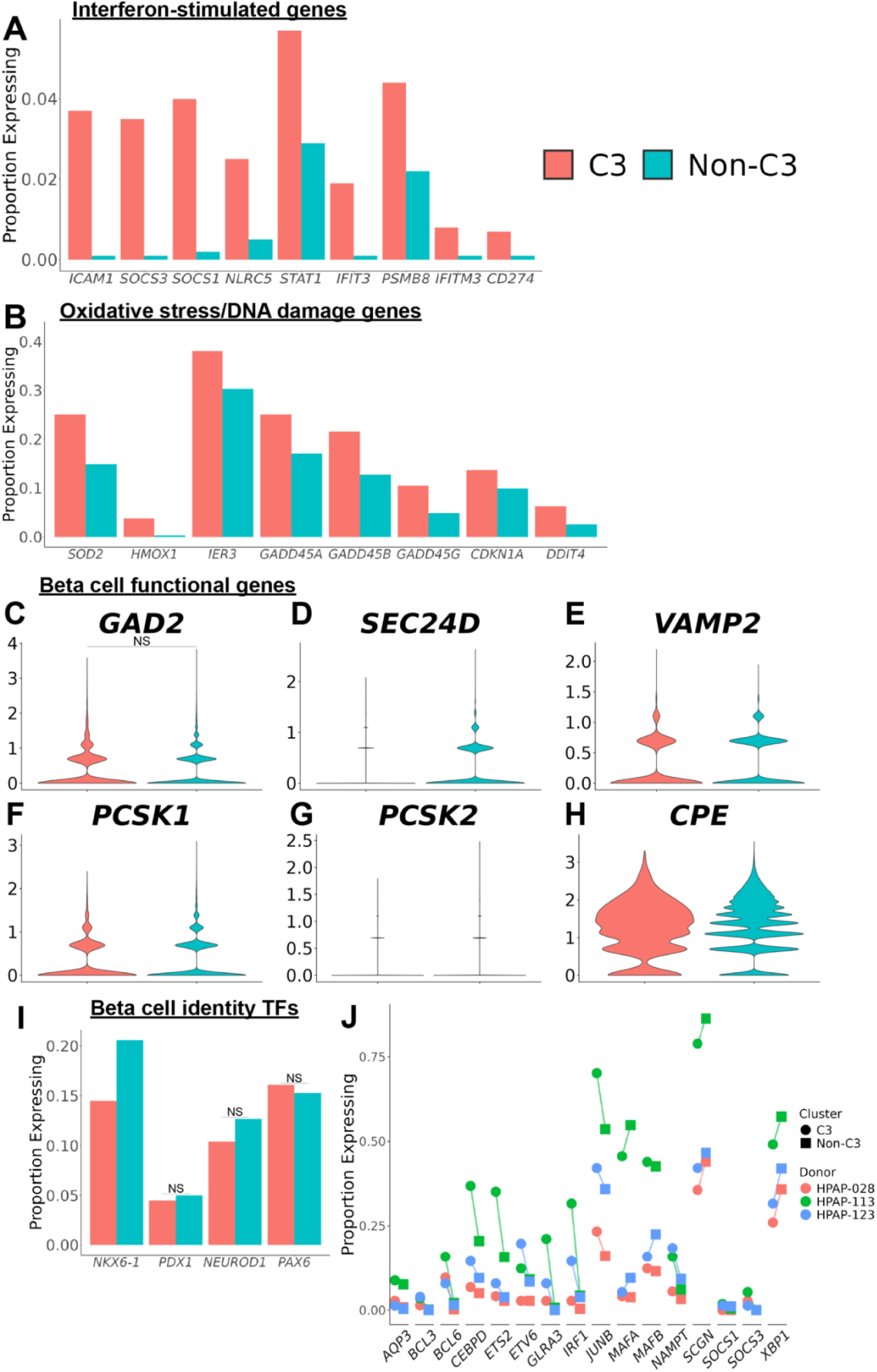
Transcriptional profile of cluster 3 beta cells. A: Frequency of beta cells expressing indicated ISGs between C3 and non-C3 beta cells. B: Frequency of beta cells expressing indicated oxidative stress/DNA damage genes between C3 and non-C3 beta cells. C-H: Violin plots showing expression of indicated transcripts for C3 and non-C3 beta cells. I: Frequency of beta cells expressing lineage-defining beta cell TFs between C3 and non-C3 beta cells. For all panels, all adjusted p values (Bonferroni) < 0.05 unless indicated by NS (not significant). P values derived from DEG testing (Methods). Full cluster 3 differential expression tabular data is available in Supplementary Table 9. J: Percent of cells expressing indicated transcripts for the 3 T1D samples with the most C3 beta cells. Trends are shown (not significance) due to low cell counts in individual donors.

**Supplementary Fig. 9:**
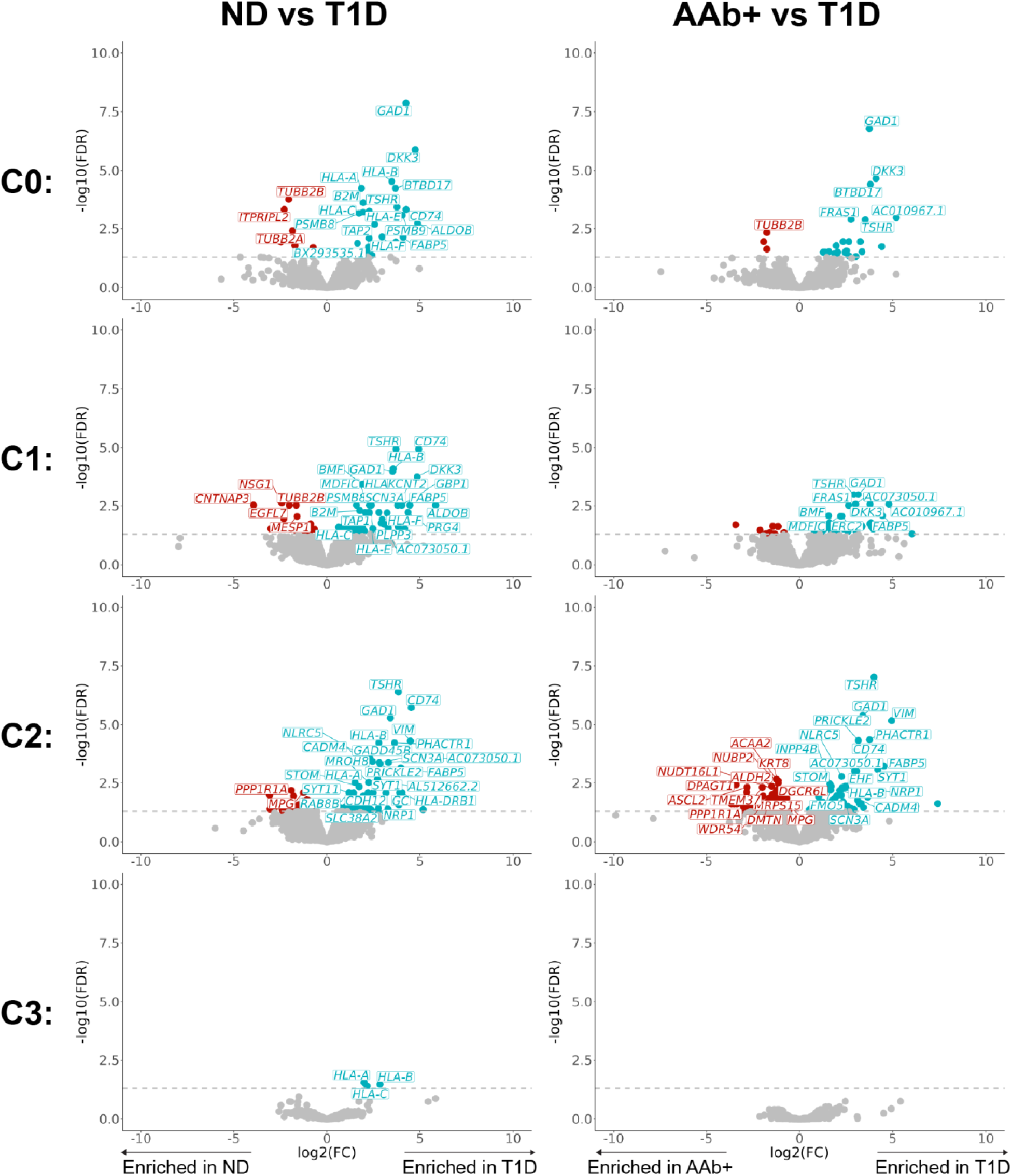
DEGs between T1D and ND/AAb+ beta cells within each beta cell subcluster. Volcano plots of DEGs between T1D and ND/AAb+ beta cells within each cluster calculated using edgeR with donor age, sex, BMI and 10X Chromium kit chemistry as covariates. A pseudobulk method was used by aggregating counts by donor. Donors with less than 5 beta cells in a particular cluster were excluded for comparisons within that cluster. BH-calculated FDR values are shown on the y axis. Dashed lines indicate FDR = 0.05.

**Supplementary Fig. 10:**
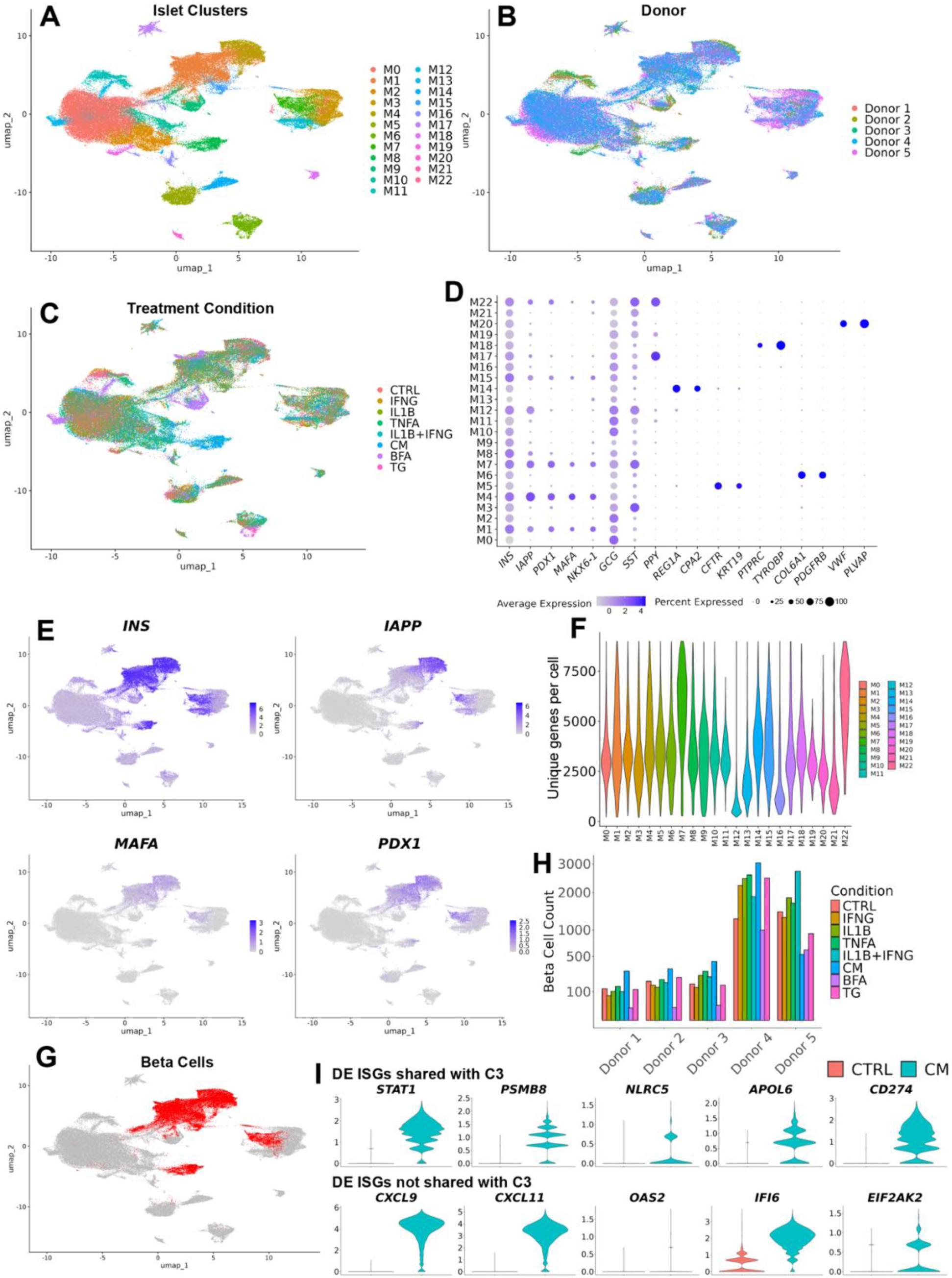
Beta cell identification and gene expression from Maestas et al. in vitro scRNAseq data. A: Integrated UMAP from reanalysis of in vitro islet scRNAseq data from Maestas et al. B-C: UMAP showing cells colored by donor (B) and treatment condition (C; CTRL: control, CM: cytokine mix with IFNG+IL1B+TNFA, BFA: Brefeldin A, TG: Thapsigargin). Treatments were applied for 48 hours. D: Dot plot of scaled islet marker gene expression within each cluster. E: Feature plots of beta cell marker transcripts. F: The number of unique genes expressed per cell within each cluster. G: UMAP showing cells identified as beta cells from Maestas et al. scRNAseq data (highlighted in red, beta cells defined as cells belonging to clusters M1, M4, M7, M8, or M15). H: Beta cell counts recovered for each sample (donor + treatment combination). I: Violin plots of in vitro cytokine-treated and control beta cells (pooled from all donors). All panels are differentially expressed with adjusted p values (Bonferroni) < 0.05. Shared ISGs are those differentially expressed in both in vivo C3 vs non-C3 beta cells (HPAP) and in vitro cytokine-treated vs control beta cells (Maestas et al.). Non-shared ISGs are those only differentially expressed by in vitro cytokine-treated vs control beta cells (Maestas et al.). Full tabular data for HPAP C3 DEGs are found in Supplementary Table 9 and for Maestas et al. in Supplementary Table 13.

**Supplementary Fig. 11:**
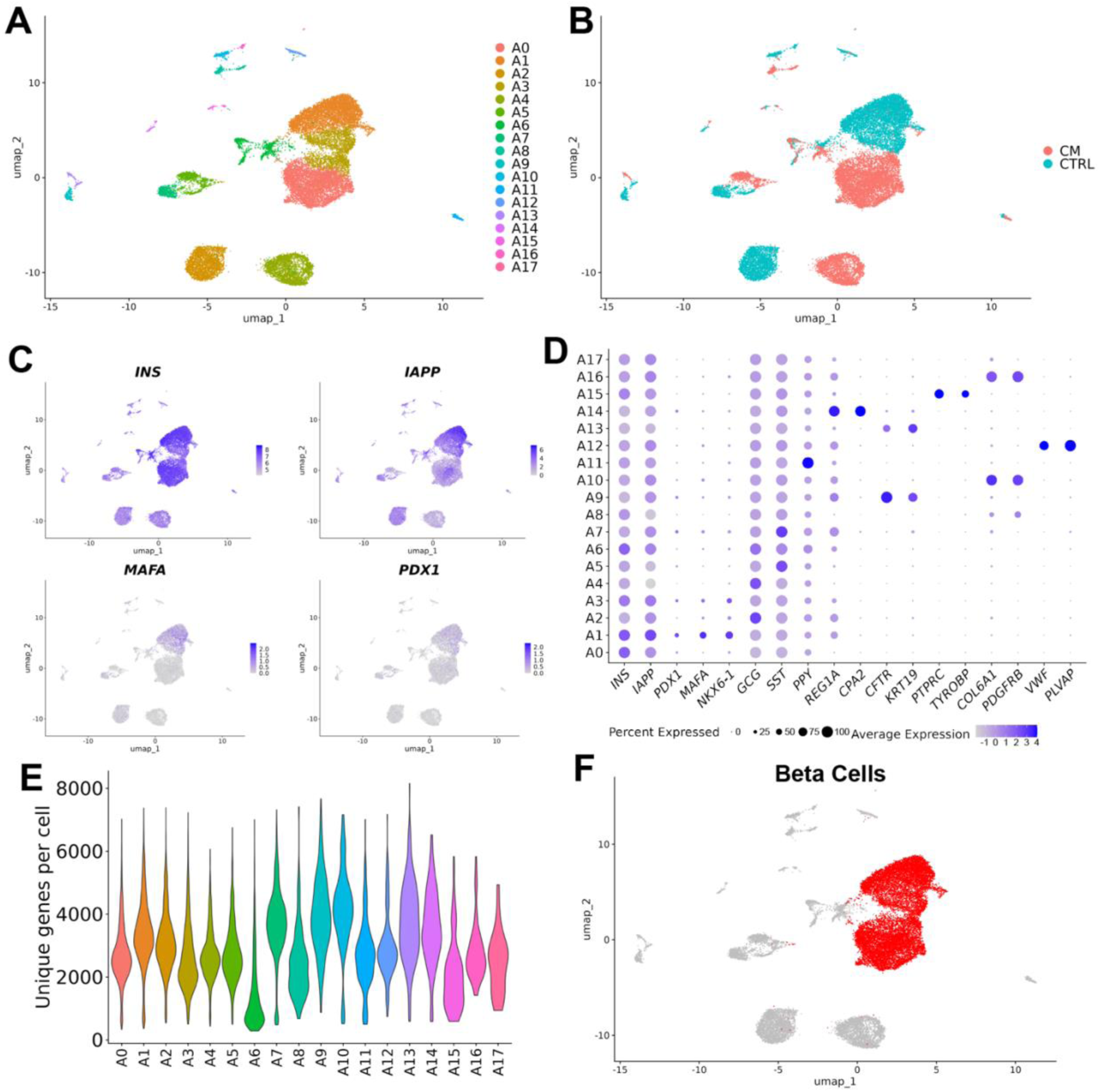
Beta cell identification from Maestas et al. in vitro snMultiome data. A: Integrated ATAC and RNA UMAP from Maestas et al. islet snMultiome data. B: UMAP showing cells colored by treatment condition (CTRL: control, CM: cytokine mix with IFNG+IL1B+TNFA; 48 hour treatment duration). C: Feature plots of beta cell marker transcripts. D: Dot plot of scaled islet marker gene expression within each cluster. E: The number of unique genes expressed per cell within each cluster. F: UMAP showing cells identified as beta cells from Maestas et al. snMultiome data (highlighted in red, beta cells defined as cells belonging to clusters A0, A1, or A3).

**Supplementary Fig. 12:**
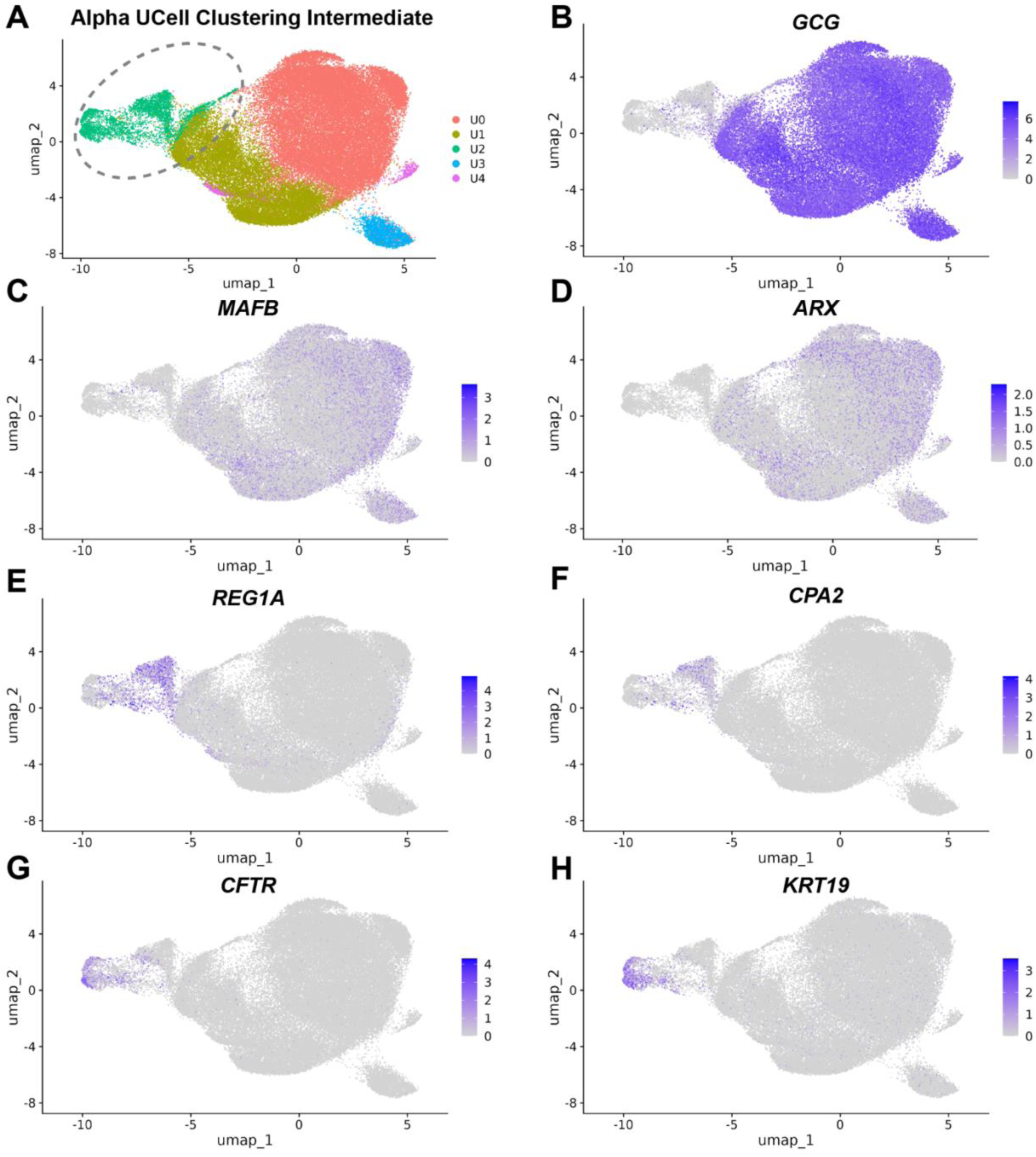
Intermediate alpha cell UCell clustering. A: Intermediate alpha cell clustering based on UCell scores (Methods). Gene sets for UCell score calculation are in Supplementary Table 3. Cells in cluster U2 (circled) were identified as likely contaminants and removed. B-H: Feature plots showing normalized expression of indicated transcripts.

**Supplementary Fig. 13:**
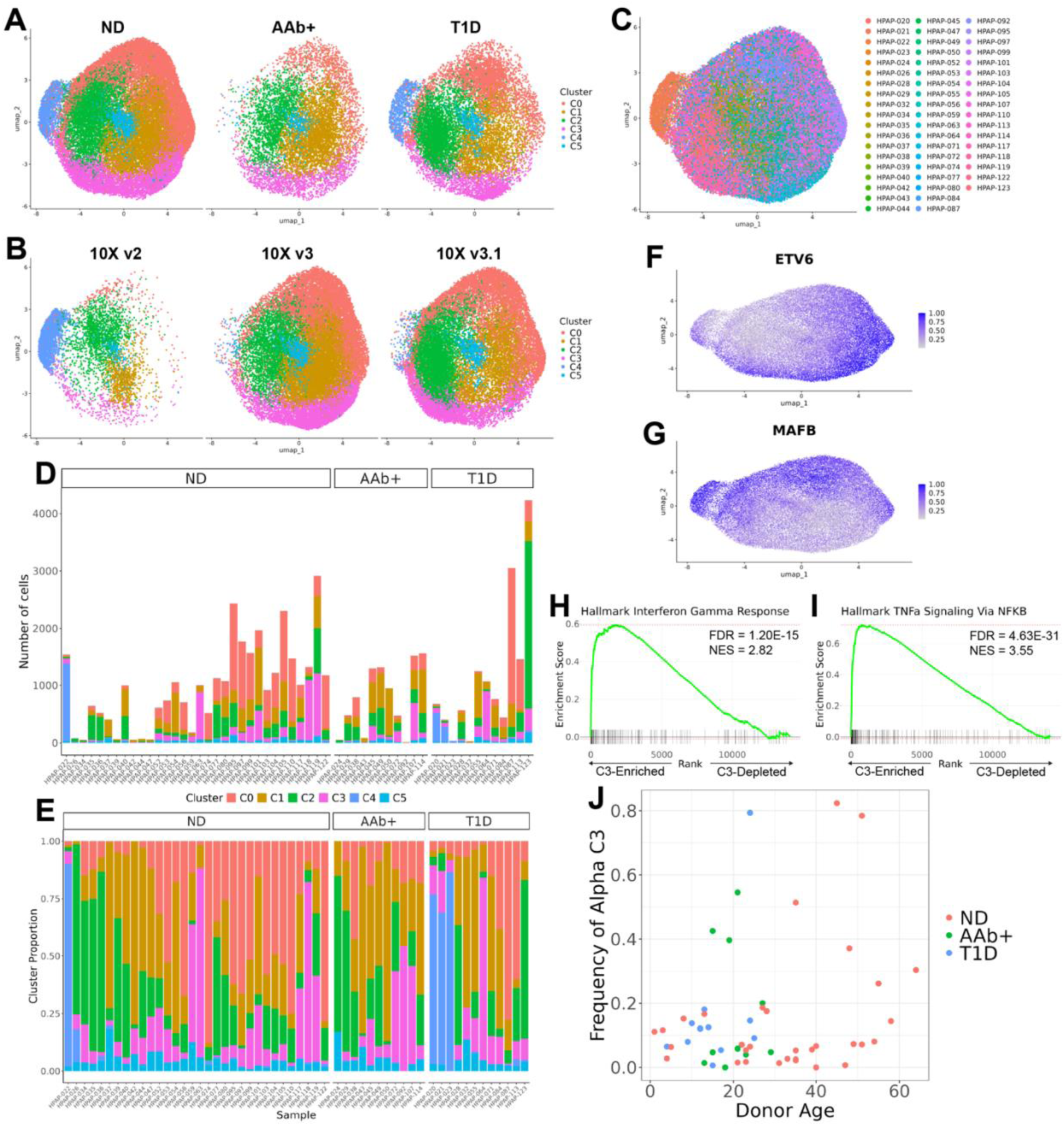
Identification of an alpha cell population homologous to cluster 3 beta cells. A-B: Alpha cell UMAP split by clinical status (A) and 10X Chromium reagent kit chemistry (B). C: Alpha cell UMAP by individual donor. D: Count of each alpha cell cluster per donor. E: Frequency of each alpha cell cluster per donor. F-G: Feature plots of rank-normalized regulon activity scores. H-I: Gene set enrichment plots for selected pathways (full list of gene sets in Supplementary Table 3). Genes were ranked by fold change in C3 alpha cells (vs non-C3 alpha cells), with filtering of lowly expressed genes (Methods, full table of GSEA output in Supplementary Table 20). BH-calculated FDR values are shown. NES: normalized enrichment score. J: Frequency of C3 alpha cells (among all alpha cells) versus donor age. Each dot is one donor, colored by clinical status.

**Supplementary Fig. 14:**
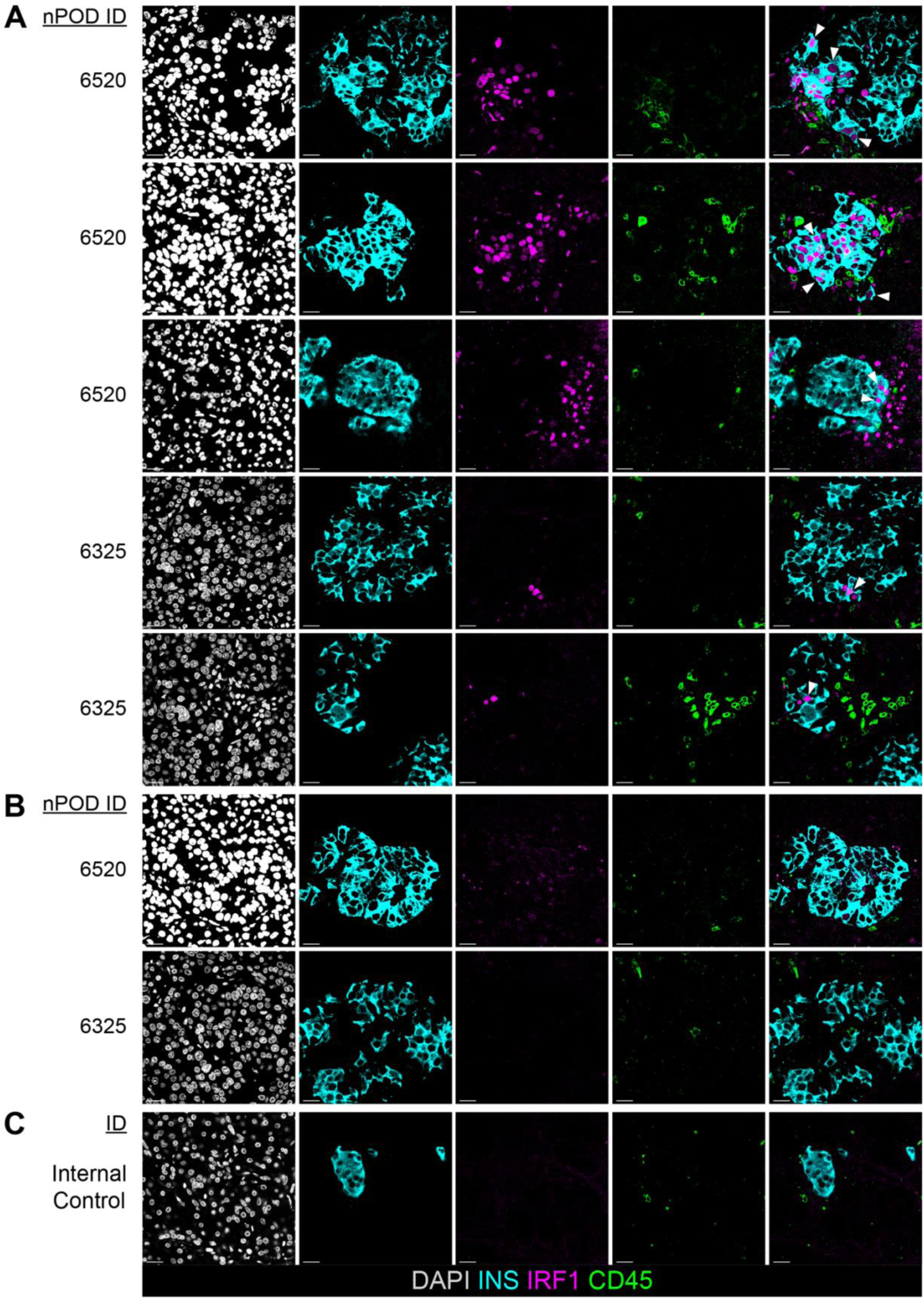
The spatial distribution of C3 beta cells in T1D islets. INS (cyan), IRF1 (magenta), CD45 (green), and a nuclear marker (DAPI; grey) jointly visualized in FFPE pancreatic tissue sections, as in Fig. 6. Single channel and composite images (far right column) are shown. Brightness and contrast were adjusted to enhance visualization. Arrows indicate C3 beta cells (INS+IRF1+ cells). Scale bars correspond to 20μm. Additional clinical information is present in Supplementary Table 21. A: T1D islets containing C3 beta cells. B: Islets lacking C3 beta cells in T1D donors with abundant C3 beta cells in other islets. C: Images from an internal ND control.

**Supplementary Table 21.** Clinical characteristics of individuals included in immunofluorescence studies.

| nPOD ID | Age | Sex | T1D Duration | Additional Information |
| --- | --- | --- | --- | --- |
| 6520 | 22 | Male | 0 years (unsuspected) |  |
| 6551 | 21 | Male | 0.6 years |  |
| 6579 | 14 | Female | 1 year |  |
| 6325 | 20 | Female | 6 years |  |
| 6590 | 16 | Female | Non-diabetic |  |
| Internal Control | 76 | Female | Non-diabetic | PDAC resection adjacent healthy tissue |

## Notes

**Conflicts of Interest:** KCH has a patent for the use of teplizumab for delay of Type 1 diabetes but no financial interest.

https://hpap.pmacs.upenn.edu

https://www.ncbi.nlm.nih.gov/geo/query/acc.cgi?acc=GSE205853

### Competing Interest Statement

Dr. John Tsang serves on the scientific advisory boards of CytoReason Inc. and Immunoscape Inc. and as the co-chief scientific officer (as an unpaid volunteer) of the Human Immunome Project. Dr Kevan Herold. has a patent for the use of teplizumab for delay of type 1 diabetes (patent US20220041720, issued February 10, 2022) but no financial interest.

### Summary of Updates

1)A new panel in Figure 2 (Fig. 2E) which examines C3 frequency by T1D disease duration, as raised by multiple reviewers. 2)A modified Discussion section which emphasizes that the C3 state promoting beta cell survival cannot be established from cross-sectional, cadaveric data, and that other interpretations cannot be excluded. This caveat was also raised by multiple reviewers. 3)A new figure (Figure 6, along with Supplementary Figure 14 and Supplementary Table 21) which uses immunofluorescence to orthogonally validate the presence of C3 beta cells in tissues from donors with T1D.

